# Discovery of novel microRNA mimic repressors of ribosome biogenesis

**DOI:** 10.1101/2023.02.17.526327

**Authors:** Carson J. Bryant, Mason A. McCool, Gabriela T. Rosado-González, Laura Abriola, Yulia V. Surovtseva, Susan J. Baserga

**Author notes:** Address correspondence to Susan J. Baserga (Molecular Biophysics and Biochemistry, PO Box 208024, 333 Cedar Street, New Haven, CT 06520-8024, USA).

## Abstract

While microRNAs and other non-coding RNAs are the next frontier of novel regulators of mammalian ribosome biogenesis (RB), a systematic exploration of microRNA-mediated RB regulation has not yet been undertaken. We carried out a high-content screen in MCF10A cells for changes in nucleolar number using a library of 2,603 mature human microRNA mimics. Following a secondary screen for nucleolar rRNA biogenesis inhibition, we identified 72 novel microRNA negative regulators of RB after stringent hit calling. Hits included 27 well-conserved microRNAs present in MirGeneDB, and were enriched for mRNA targets encoding proteins with nucleolar localization or functions in cell cycle regulation. Rigorous selection and validation of a subset of 15 microRNA hits unexpectedly revealed that most of them caused dysregulated pre-rRNA processing, elucidating a novel role for microRNAs in RB regulation. Almost all hits impaired global protein synthesis and upregulated *CDKN1A* (*p21*) levels, while causing diverse effects on RNA Polymerase 1 (RNAP1) transcription and TP53 protein levels. We discovered that the MIR-28 siblings, hsa-miR-28-5p and hsa-miR-708-5p, directly and potently target the ribosomal protein mRNA *RPS28* via tandem primate-specific 3’ UTR binding sites, causing a severe pre-18S pre-rRNA processing defect. Our work illuminates novel microRNA attenuators of RB, forging a promising new path for microRNA mimic chemotherapeutics.

**Graphical Abstract:** 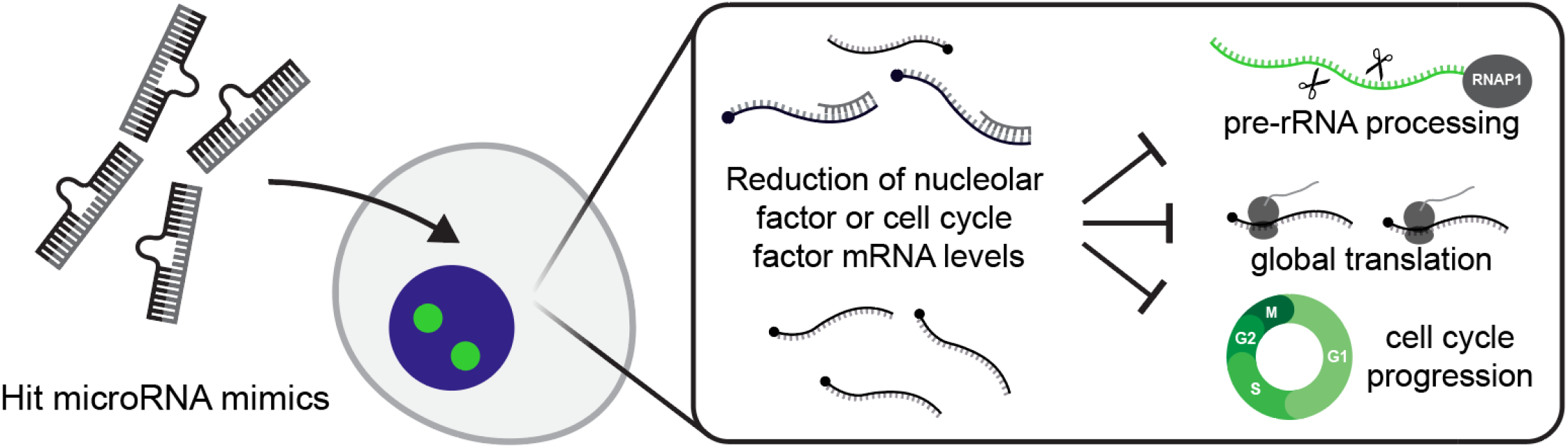

## Introduction

Ribosome biogenesis (RB) is the complex, essential process by which mature small and large ribosomal subunits are produced in all living organisms. Eukaryotes partition many RB steps into the nucleolus, a phase-separated membraneless organelle within the enveloped nucleus (1–3). In human cells, three of the four mature ribosomal RNAs (rRNAs), the 18S, 5.8S, and 28S rRNAs, are synthesized in the nucleolus as components of the polycistronic 47S primary pre-rRNA precursor transcript from tandem ribosomal DNA (rDNA) repeats by RNA Polymerase 1 (RNAP1) (4). The 5S rRNA is separately transcribed in the nucleus by RNA Polymerase 3 (RNAP3) (5,6). A myriad ribosome assembly factors (AFs) execute endo- and exonucleolytic pre-rRNA processing and modification events to liberate the mature rRNAs from the 47S transcript, forming the small 40 large 60S ribosomal subunits (7–10). AFs also facilitate the binding of structurally-constitutive ribosomal proteins (RPs) and the folding of the maturing subunits at the macromolecular scale (11–16). Defects in RB can trigger the nucleolar stress response during which labile members of the 5S RNP including RPL5 (uL18) or RPL11 (uL5) bind and sequester the TP53-specific E3 ligase MDM2, effectively stabilizing TP53 levels and leading to *CDKN1A* (*p21*) induction, cell cycle arrest, and apoptosis (17–19). At the organismal level, nucleolar stress resulting from RB defects can cause a class of rare human diseases called ribosomopathies (19–22). Furthermore, cancer initiation and progression are strongly linked to aberrant RB (23–29).

MicroRNAs comprise a class of non-coding (nc)RNAs of approximately 22 nt which can base pair with messenger (m)RNAs to post-transcriptionally reduce transcript stability or translation efficiency, acting as “sculptors of the transcriptome” to fine-tune gene expression (30–32). Like RB, microRNAs play critical roles in mediating human development, health, and disease including cancer (33,34). How microRNAs regulate RB has yet to be explored systematically at the experimental level, although some intriguing links between them have been uncovered. A handful of microRNAs have been experimentally demonstrated to affect RB subprocesses including RNAP1 transcription, 60S assembly, and RP gene transcription (35). Consistent with this, the microRNA biogenesis factors Drosha and Dicer are required for 28S and 5.8S maturation (36). AGO2, the microRNA-binding component of the active RISC complex (32), has been found in the nucleolus (37) along with several microRNAs (38–40), though their precise biological function there remains unclear. Computational analysis has implicated microRNA-mediated control of RPs as a key potentiator of RB activity and disease progression (41), thereby necessitating additional *in vivo* experiments. Software packages have made some inroads towards accurate prediction of microRNA targets (42,43) or functions (44) although abundant false positives limit their utility (41,45). The limited amount of direct experimental evidence that microRNAs are involved in RB constitutes a significant gap in our understanding of the layers of regulation of nucleolar function in human cells.

To date, no holistic, unbiased discovery campaign for microRNAs functioning in RB regulation has been conducted, and the full complement of microRNAs affecting RB remains poorly understood. We previously hypothesized that microRNAs may be a key underappreciated conduit linking biochemical RB defects to the pathogenesis of diseases like ribosomopathies and cancer (35). To discover novel microRNAs negatively regulating RB, we conducted our previously-established high-content screen for changes in nucleolar number following microRNA mimic overexpression in human MCF10A cells. High-throughput screens using microRNA mimics have previously revealed mechanistic insight into microRNA-mediated regulation dynamics during cellular proliferation and signaling (46,47), cardiac regeneration (48), viral infection (49), and cancer (50,51).

Here, we identify 72 high-confidence mature human microRNA hits that disrupt RB, which are enriched for mRNA targets involved in cell cycle regulation, cellular proliferation, and localization within the nucleolus. Twenty-seven (27) of these hits are validated, conserved microRNAs present in the MirGeneDB database (52). We validate the roles of a subset of 15 hits in RB subprocesses including pre-rRNA transcription, pre-rRNA processing, and global protein synthesis. For the first time, we directly define the abilities of 12 microRNA mimics to inhibit pre-rRNA processing. Our work reveals that the MIR-28 family members, hsa-miR-28-5p and hsa-miR-708-5p, are strong inhibitors of pre-18S pre-rRNA processing by way of direct and potent downregulation of the levels of the ribosomal protein mRNA, *RPS28*. Our screen’s results underscore the broad potential of microRNAs to dysregulate RB, and raise new questions regarding the extent to which microRNAs may connect RB and disease.

## Materials and Methods

### Cell lines and culture conditions

Human MCF10A breast epithelial cells (ATCC CRL-10317) were cultured in DMEM/F-12 (Gibco 11330032) with 5% horse serum (Gibco 16050122), 10 µg/mL insulin (MilliporeSigma I1882), 0.5 µg/mL hydrocortisone (MilliporeSigma H0135), 20 ng/mL epidermal growth factor (Peprotech AF-100- 15), and 100 ng/mL cholera toxin (MilliporeSigma C8052). hTERT RPE-1 cells (ATCC CRL-4000) were cultured in DMEM/F-12 (Gibco 11330032) with 10% fetal bovine serum (Gibco 16050122) and 1% Pen-Strep (Gibco 15140122). HeLa cells (ATCC CCL-2) were cultured in in DMEM (Gibco 11965092) with 10% fetal bovine serum (Gibco 16050122). Cells were incubated at 37 °C in a humidified atmosphere with 5% CO2. HEK 293 Flp-In T-REx cells (Invitrogen R75007) were a generous gift from P. Gallagher, Yale School of Medicine, New Haven, CT, and were grown in DMEM (Gibco 11965092) with 10% fetal bovine serum (Gibco 16050122) and 15 μg/mL blasticidin S (Alfa Aesar J67216XF).

### Chemical reagents

BMH-21 (Sigma-Aldrich SML1183; CAS 896705-16-1) was diluted to a working stock concentration of 50 μM in DMSO for direct dosing of cells in 384-well plates.

### RNAi depletion and microRNA expression by reverse-transfection

RNAi depletion by reverse-transfection was conducted in MCF10A cells, hTERT RPE-1 cells, and HeLa cells as previously reported (53–55). Briefly, cells were reverse-transfected into an arrayed 384- well plate library containing small interfering RNA (siRNA) or miRIDIAN microRNA mimic constructs (Horizon Discovery, **Supplementary Tables 1-7**) using Opti-MEM (Gibco 31985070) and Lipofectamine RNAiMAX transfection reagent (Invitrogen 13778150). Assay-ready plates containing 10 µL of 100 nM microRNA mimics resuspended in 1X siRNA buffer (Horizon Discovery B-002000- UB-100) were prepared from master library 384-well plates (Horizon Discovery, 0.1 nmol scale) and stored at -80 C. Plates were prepared with control siRNAs (siNT, siNOL11, siKIF11, or siPOLR1A) for reverse-transfection at a final 20 nM siRNA/microRNA mimic concentration as described (55), at a seeding density of 3000 MCF10A cells/well.

### 5-ethynyl uridine labeling; staining and high-content imaging

5-ethynyl uridine (5-EU; ClickChemistryTools 1261-100, CAS 69075-42-9) was used to label cells at a 1 mM final concentration. Staining, click chemistry, and high-content imaging were performed as previously described (55).

### CellProfiler pipeline and data analysis

Image analysis and data processing were conducted using a custom pipeline for CellProfiler 3.1.9 as previously described (53–57). Strictly-standardized mean difference (SSMD) values were calculated from plate-adjusted one-nucleolus or 5+ nucleoli percent effect values using the uniformly minimal variance unbiased estimate (UMVUE), equation A5 in (58). Data from the primary or secondary screen were averaged in JMP (version 17.0, SAS Institute, Cary, NC 1989-2023) and graphed with JMP or GraphPad Prism 8 (GraphPad Software).

### MCF10A RNA expression dataset

Deposited reads from four RNAseq experiments from MCF10A cells quantifying transcript levels without treatment (negative control conditions; see table below) were re-analyzed using Partek Flow. Reads were aligned to the hg38 genome with HISAT2 2.1.0 and quantified using the Ensembl Transcripts version 99 annotation with the Partek E/M algorithm module. Normalized transcripts per million (zTPM) for genes in each experiment was calculated in R as described (59). For each dataset, a given gene was categorized as expressed if its zTPM was greater than -3, as described in (59) (**Supplementary Table 8**).

| BioProject accession | GEO samples used | Reference |
| --- | --- | --- |
| PRJNA290557 | GSM1829628 | (60) |
| PRJNA384982 | GSM2593351, GSM2593352, GSM2593353 | (61) |
| PRJNA530983 | GSM3711368, GSM3711369 | N/A |
| PRJNA647393 | GSM4667014, GSM4667015, GSM4667016 | (62) |

### Nucleolar protein metadatabase

Three nucleolar protein databases were merged to create a nucleolar protein metadatabase. The Human Protein Atlas subcellular localization database (v. 20.0) (63, proteinatlas.org) containing 1350 unique nucleolar proteins was retrieved, and HGNC gene names were updated based on Ensembl Gene ID using BioMart. Proteins labeled with the GO terms “Nucleoli”, “Nucleoli fibrillar center”, or “Nucleoli rim” were considered to be nucleolar. The T-cell nucleolar proteome (64) containing 880 unique nucleolar proteins was retrieved, and HGNC gene names were updated based on NCBI Entrez Gene ID using BioMart. An archived copy of NOPdb 3.0 (65) containing 2242 unique nucleolar proteins was retrieved, and HGNC gene names were updated based on International Protein Index (IPI) ID using the latest IPI database release (v. 3.87) and BioMart. The three databases were merged on updated HGNC name, resulting in a metadatabase of 3490 unique nucleolar proteins (**Supplementary Table 9**).

### Bioinformatic target enrichment analysis

TarBase 8 (66) was utilized to identify experimentally-validated microRNA:mRNA interactions. Genes were filtered for the *Homo sapiens* species, for true positive microRNA:mRNA interactions, and for interactions where the microRNA caused reduced levels of the mRNA. Next, genes were annotated with zTPM expression data from the MCF10A expression dataset (see above), for nucleolar localization using the nucleolar protein metadatabase (see above), and 89 cytosolic RP genes were labeled based on HGNC gene groups (35 “S ribosomal proteins”, HGNC gene group 728; 54 “L ribosomal proteins”, HGNC gene group 729). The number of genes targeted by each microRNA was calculated in JMP, and confirmed hit microRNAs were labeled. Subset tabulations were also carried out to determine the number of genes coding for nucleolar proteins targeted by each microRNA, and the number of genes coding for RPs targeted by each microRNA. Mean and median values for each category were calculated with JMP for hit microRNAs and non-hit microRNAs. Conversely, the number of microRNAs targeting each gene was also calculated with JMP, and all 262 genes targeted by 5 or more microRNA hits were analyzed for enrichment using Enrichr (67–69) (**Supplementary Table 10**). For the subset of 24 evolutionarily-conserved MirGeneDB hits with validated targets from the primary screen, 135 genes targeted by 5 or more hits were analyzed for enrichment using Enrichr (**Supplementary Table 10**). To analyze the evolutionary conservation of the MIR-28 binding sites in the *RPS28* 3’ UTR, indicated species from the Multiz 100 Vertebrate Species Alignment and Conservation track in the UCSC Genome Browser were visualized (70,71).

### RNA isolation following RNAi transfection

MCF10A cells or hTERT RPE-1 cells were seeded at 100,000 cells per well in 2 mL of medium in 6- well plates and incubated at 37 °C for 24 h. For the MIR-28 dose response, hTERT RPE-1 cells were seeded at 50,000 cells per well in 1 mL of medium in 12-well plates and incubated as above. Cells were transfected with siRNAs or microRNA mimics at 30 nM or as otherwise indicated using Lipofectamine RNAiMAX (Invitrogen 13778-150) and Opti-MEM (Gibco 31985070) per manufacturer’s instructions for 72 h. Cells were washed with 1X PBS, then collected with 1 mL of TRIzol reagent (Invitrogen 15596026). Total RNA was purified following the manufacturer’s protocol.

### PolyA+ RNAseq following overexpression of hsa-miR-28-5p or hsa-miR-708-5p

MCF10A cells or hTERT RPE-1 cells were treated with siNT, hsa-miR-28-5p, or hsa-miR-708-5p, and RNA was isolated as above. One µg of total RNA was resuspended in nuclease-free H_2_O and submitted to the Yale Center for Genomic Analysis (West Haven, CT) for polyA+ library preparation and sequencing. All samples had an RNA Integrity Number (RIN) of at least 9.6. 30-50 million 100 bp paired-end reads were collected per sample using a NovaSeq 6000 Sequencing System (Illumina).

RNAseq was conducted in biological triplicate. Partek Flow was used to process, align, and quantify reads. Reads were trimmed of adapters, then aligned to the hg38 genome with HISAT2 2.1.0. Reads were quantified using the RefSeq Transcripts version 93 annotation with the Partek E/M algorithm module. Differential expression analysis was conducted with DESeq2 (72). Raw reads are available on NCBI under BioProject accession PRJNA919164, and processed data are summarized in GEO accession GSE242754 and **Supplementary Tables 11-12**.

### Analysis of RNA transcript levels by RT-qPCR

MCF10A cells or hTERT RPE-1 cells were treated with control siRNAs or microRNA mimics, and RNA was isolated as above. cDNA was synthesized from 1 µg total input RNA using iScript^TM^ gDNA Clear cDNA Synthesis Kit (Bio-Rad 1725035). In a qPCR plate (Bio-Rad MLL9601), 1 µL of cDNA was dispensed, followed by 19 µL of a qPCR master mix containing iTaq Universal SYBR Green Supermix (Bio-Rad 1725121), 500 nM forward primer, 500 nM reverse primer, and water. Technical duplicates were assayed for each biological replicate. The plate was briefly centrifuged and assayed using a Bio-Rad CFX96 Touch Real-Time PCR Detection System. Amplification parameters were: initial denaturation 95 °C for 30 s; 40 cycles 95 °C denaturation for 15 s, 60 °C annealing and extension for 30 s. Melt curve analysis parameters were: 60 °C to 94.8 °C in 0.3 °C increment. Data analysis was completed using the comparative C_T_ method (ΔΔC_T_) using 7SL RNA as an internal loading control.

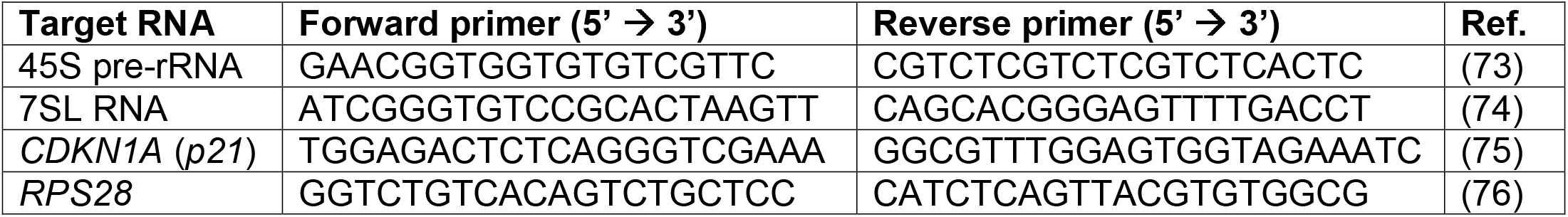

### Analysis of mature rRNAs

MCF10A cells, hTERT RPE-1 cells, or HEK 293 Flp-In cells were treated with control siRNAs or microRNA mimics, and RNA was isolated as above. One μg of total RNA was resuspended in nuclease-free H_2_O and submitted to the Yale Center for Genomic Analysis for electropherogram analysis. Each experiment was conducted with either a Bioanalyzer 2100 or a Fragment Analyzer 5300 (Agilent). Mature rRNA ratios and mature rRNA relative peak areas were taken from the output reports. The data were graphed and analyzed by ANOVA followed by Holm-Šídák post-hoc testing in GraphPad Prism.

### Northern blot analysis of pre-rRNA processing

MCF10A cells were treated with siRNAs or microRNA mimics, and RNA was isolated as above. Northern blots were performed using 3 μg of total RNA as published (53,54), and were performed in at least biological triplicate. Blots were quantified with Image Lab 6.0.1 (Bio-Rad). RAMP [Ratio Analysis of Multiple Precursors, (77)] ratios were calculated in Microsoft Excel, and heatmaps were made using the mean log_2_ RAMP ratio for each treatment relative to siNT in GraphPad Prism. The following DNA oligonucleotides were radiolabeled for blotting:

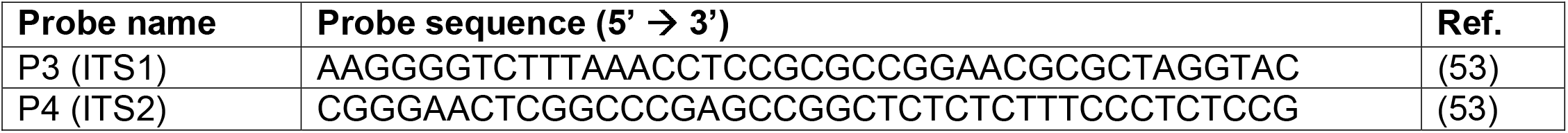

### Dual-luciferase assay for RNAP1 promoter activity

MCF10A cells were seeded at 30,000 cells per well in 1 mL of medium in 12-well plates and incubated at 37 °C for 24 h. Cells were transfected with 30 nM siRNAs or microRNA mimics as above for 48 h. Cells were then transfected with 1 μg of pHrD-IRES-Fluc (Addgene plasmid #194250) and 1 ng of CMV-Rluc reporter plasmids (78) using Lipofectamine 3000 (Invitrogen L3000015).

Concurrently, cells were treated with 3.5 μL of DMSO vehicle or 300 μM BMH-21, to achieve a final concentration of 1 μM BMH-21. After another 24 h, treated cells were washed once with 1X PBS and lysed with 250 μL of 1X Passive Lysis Buffer (Promega E1941) at room temperature for at least 30 minutes. In a solid white 96-well plate (Greiner Bio-One 655074), 20 μL of lysate from each sample was dispensed into a well. Samples were assayed using a Promega GloMax plate reader with dual injectors, using the Dual-Luciferase Reporter Assay System (Promega E1910) per manufacturer’s instructions. Sixty μL of LAR II or Stop & Glo substrate were injected with a 2 s delay and a 10 s read time. Data were analyzed by calculating the Fluc/Rluc ratio for each well, then normalizing to the Fluc/Rluc ratio for siNT. Data import was carried out in Microsoft Excel, calculations were performed in JMP, and normalized data were graphed and analyzed by ANOVA followed by Holm-Šídák post-hoc testing in GraphPad Prism.

### Protein isolation, SDS-PAGE analysis, and immunoblotting

MCF10A cells were seeded at 100,000 cells per well in 2 mL of medium in 6-well plates and incubated at 37 °C for 24 h. Cells were transfected with a final concentration of 30 nM siRNAs or microRNA mimics in a total volume of 2250 μL as above for 72 h. Following treatment, cells were washed twice with cold 1X PBS, manually dislodged using cell scrapers (Falcon 353085), collected in 1 mL of cold 1X PBS, centrifuged at 1100 RCF for 5 minutes at 4 °C, then lysed in AZ lysis buffer (50 mM Tris pH 7.5, 250 mM NaCl, 1% Igepal, 0.1% SDS, 5 mM EDTA pH 8.0) with 1X complete protease inhibitors (cOmplete Protease Inhibitor Cocktail, Roche 11697498001) by vortexing. Cell debris was pelleted at 21000 RCF for 15 minutes. A Bradford assay was used to determine supernatant total protein concentration. Protein aliquots were made using 5X Laemmli buffer, then boiled at 95°C for 3 minutes and loaded onto a gel or frozen at -20°C. Handcast SDS-PAGE gels (8%, 10%, 15%, or 4-18% gradient) containing 0.5% (v/v) trichloroethanol (Acros Organics 139441000) for stain-free imaging (79) were used to separate total protein at 110 V for 2 h. Total protein was imaged using the ChemiDoc stain-free imaging protocol (Bio-Rad) to ensure even loading at the gel stage. Gels were UV-activated for 5 min in the ChemiDoc. Following membrane transfer with the Trans-Blot Turbo system (Bio-Rad), blots were imaged again for total protein; these images are presented and quantified as blot loading controls. Immunoblotting was carried out using 5% (w/v) Omniblok dry milk (American Bio AB10109) in 1X PBST (1X PBS containing 5% (v/v) Tween) with primary antibodies listed below followed by 1:5000 peroxidase-linked anti-mouse or anti-rabbit IgG (Amersham NXA931 or NA934) as appropriate. Blots were developed using low- or high-sensitivity ECL reagent (Millipore WBKLS0500, Thermo Scientific 34094) for 5 minutes, then dried and imaged with the ChemiDoc. Images from the ChemiDoc were quantified using Image Lab 6.0.1 (Bio-Rad).

Data were graphed and analyzed by ANOVA followed by Holm-Šídák post-hoc testing in GraphPad Prism.

| **Target molecule** | **Primary antibody manufacturer and catalog number** | **Dilution** |
| --- | --- | --- |
| Puromycin | Kerafast EQ0001 (clone 3RH11) | 1:5000 |
| RPS28 | Invitrogen PA5-45721 | 1:500 |
| TP53 | Santa Cruz Biotechnology sc-126 HRP (clone DO-1) | 1:5000 |

### Puromycin incorporation for SUnSET global translation assay

After the 72 h transfection period in 2250 μL in six-well plates, MCF10A cells were treated with an additional 750 μL of medium containing 3 μM puromycin (Mirus Bio 5940), achieving a final concentration of 1 μM (0.5 μg/mL) puromycin and final volume of 3 mL (53,54,78,80). Cells were incubated 1 h at 37 °C, then washed with cold 1X PBS. Protein was isolated and analyzed as above.

### Identification of putative microRNA binding sites

Candidate microRNA binding sites in transcripts coding for nucleolar proteins were identified using databases including TarBase 8 (66), TargetScan 7.2 (81), miRWalk 3 (82), and DIANA microT-CDS 2023 (83). Binding sites were computationally tested using the bimolecular DuplexFold algorithm on the RNAStructure Web Server (84). The sequence of either hsa-miR-28-5p or hsa-miR-708-5p was tested for *in silico* binding to 70 bp regions of target transcripts containing putative microRNA binding sites. Transcript regions bearing a scrambled binding site were also tested for binding by either mature microRNA sequence. Computed binding energies from DuplexFold are reported in this manuscript. The BLAST algorithm (85) was used to search the 45 pre-rRNA transcript (NR_046235.3) for potential binding sites for the 72 microRNA hits. BLAST matches were filtered for antisense (+/-) pairings starting on microRNA nucleotide 1 or 2 with 0 gaps to select potential canonical seed binding sites in the 45S pre-rRNA precursor (**Supplementary Table 13**).

### Molecular cloning of psiCHECK-2 plasmids for microRNA UTR assays

The psiCHECK-2 plasmid was acquired as a gift from P. Pawlica and J. A. Steitz (Yale University)

(86). Transcriptomic regions of approximately 200 bp containing putative microRNA binding sites were cloned from MCF10A genomic DNA, which was isolated using the DNeasy Blood & Tissue Kit (Qiagen 69504). Primers for generating XhoI-NotI amplicons were designed using Geneious 8.1.9 (Biomatters Ltd.) (**Supplementary Table 14**). Amplicons were restriction cloned into psiCHECK-2. Target WT seed sequences were scrambled by site-directed mutagenesis overlap cloning (87). Plasmids were verified by Sanger sequencing (GENEWIZ, Inc./Azenta Life Sciences) and are available on Addgene (plasmid #203157 and #203158).

### MicroRNA UTR assays testing for direct interaction of microRNA mimics with putative mRNA targets

MCF10A cells were seeded at 40,000 cells per well in 1 mL of medium in 24-well plates and incubated at 37 °C for 24 h. Cells were co-transfected for 24 h with 30 nM siRNA or microRNA mimic, 10 ng of psiCHECK-2 plasmid, and 1 μg of salmon sperm carrier DNA using Lipofectamine 3000 (Invitrogen L3000015) according to manufacturer protocol. Following treatment, cells were washed with 1 mL of 1X PBS and incubated in 100 μL of 1X Passive Lysis Buffer (Promega E1941) at room temperature for at least 30 minutes. The dual-luciferase assay was carried out as detailed above.

Data were analyzed by calculating the Rluc/Fluc ratio for each well, then normalizing to the Rluc/Fluc ratio for siNT. Data import was carried out in Microsoft Excel, calculations were performed in JMP, and normalized data were graphed and analyzed by ANOVA followed by Holm-Šídák post-hoc testing in GraphPad Prism.

### HEK 293 cell line engineering and MIR-28/*RPS28* rescue experiment

HEK 293 Flp-In cells were genomically engineered as described in (88), using pcDNA5 FRT/TO vectors with no insert (empty vector, EV) or a FLAG-tagged insert of the *RPS28* CDS cloned from pDONR221-RPS28 [HsCD00043374-RPS28, DNASU (89)]. Primers for generating FLAG-GGGGS-tagged BamHI-EcoRV RPS28 amplicons were designed using Geneious 8.1.9 (Biomatters Ltd.) (**Supplementary Table 14**). The FLAG-tagged RPS28 plasmid is available on Addgene (plasmid #203156). Polyclonal transformants were isolated following 200 µg/mL Hygromycin B selection (Gibco 10687010). Briefly, 75,000 HEK 293 Flp-In cells were seeded in 1 mL of medium in 12-well plates and incubated at 37 °C for 24 h. Cells were then transfected with siNT or MIR-28 microRNA mimics at 30 nM using RNAiMAX in a total transfection volume of 50 µL and were simultaneously induced with 1 µg/mL tetracycline (MilliporeSigma T7660) as indicated. Treated cells were incubated for 3 days, after which RNA was harvested using TRIzol as above. Electropherograms were collected to quantify the 28S/18S ratio using 1 µg of total RNA for each sample as above.

### miR-eCLIP analysis for identifying direct targets of MIR-28 family members

miR-eCLIP was performed as detailed in (90). Briefly, 3.5 million MCF10A cells were seeded into 15 cm tissue culture dishes and incubated for 48 h in 10 mL medium. Cells were transfected with either hsa-miR-28-5p or hsa-miR-708-5p microRNA mimics at 30 nM using RNAiMAX (see above). After 27 h, cells were washed with 15 mL 1X PBS, covered with 5 mL 1X PBS, UV crosslinked with 254 nm light at 400 mJ/cm^2^, and collected by scraping. Approximately 10 million crosslinked cells from each microRNA mimic treatment were pooled and processed for miR-eCLIP sequencing and data analysis (Eclipse Bioinnovations, San Diego, CA). Raw reads for the AGO2 immunoprecipitation (IP) and the size-matched input are available at NCBI under BioProject accession PRJNA923105. Processed data are available in GEO accession GSE242755 and **Supplementary Table 15**. Cumulative empirical distribution plots of RNAseq differential expression data for MIR-28 targets or non-targets were graphed and analyzed in R.

### Statistical testing

Statistical tests are outlined above and in the Figure Legends. Biological replicates are shown in the figures and sample sizes are noted in the Figure Legends. Statistical tests were conducted in JMP or GraphPad Prism 8. Unless otherwise stated in the Figure Legends, tests were conducted using siNT as the comparator, and *p* value magnitude is represented as **\***, *p* < 0.05; **\*\***, *p* < 0.01; **\*\*\***, *p* < 0.001.

## Results

### A high throughput phenotypic screen for altered nucleolar number identifies 71 novel microRNAs that negatively regulate ribosome biogenesis

Following on the success of our previous nucleolar number-based screens for novel RB regulators (53,54), we hypothesized that microRNAs could also be functioning as nodes of control for RB. To discover novel microRNA negative regulators of ribosome biogenesis, we screened an arrayed library of 2603 human mature microRNA mimics (Dharmacon/Horizon Discovery) for their ability to alter nucleolar number 72 h after transfection into human MCF10A cells (**Figure 1A**). While most MCF10A cells treated with a negative control non-targeting siRNA (siNT) display between two and four nucleoli, cells depleted of the tUTP NOL11 (siNOL11) have an increased probability of having one nucleolus per nucleus (“one-nucleolus phenotype”) (53,91), and those depleted of the mitotic kinesin KIF11 (siKIF11) have an increased probability of having five or more nucleoli per nucleus (“5+ nucleoli phenotype”) (54) (**Figure 1B**, siNT, siNOL11, or siKIF11 panels). Using these siRNAs as controls, we employed our established high-throughput nucleolar number screening platform to count nucleolar number after overexpression of microRNA mimics by using the CellProfiler software to segment and enumerate FBL-stained nucleoli on a per-nucleus basis from images captured by an automated microscope.

**Figure 1.**
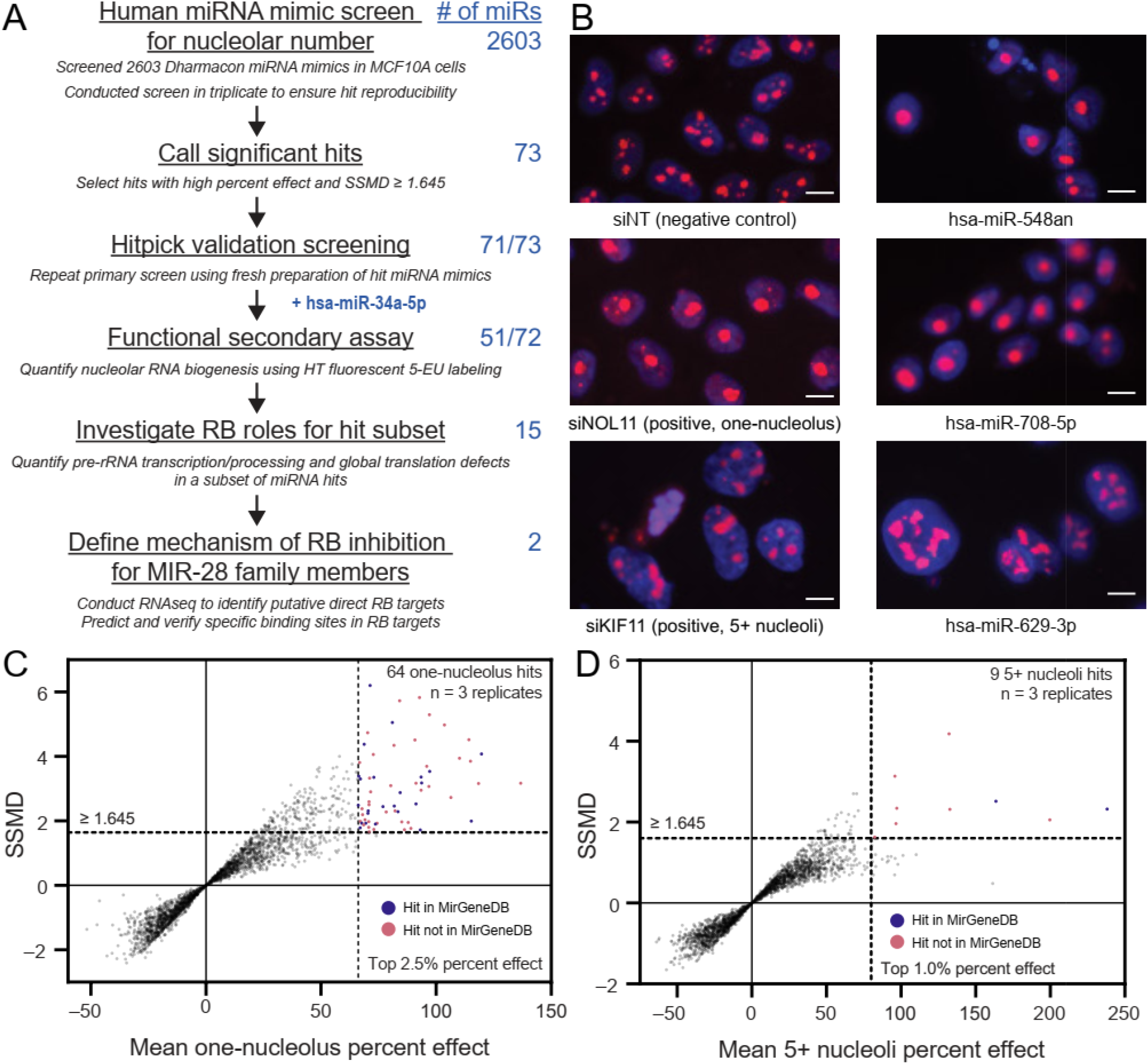
A screen for changes in nucleolar number reveals 71 novel microRNA mimic negative regulators of RB. (A) Screening campaign pipeline. MCF10A cells were reverse-transfected into a library of 2,603 mature human microRNA mimics, then fixed and stained for DNA and FBL after 72 h in biological triplicate. The number of nucleoli was calculated using CellProfiler. Hits were called based on a decrease or increase in nucleolar number, respectively termed the one-nucleolus or 5+ nucleoli phenotypes. The primary screen identified 73 high-confidence hits, 71 of which passed hitpick validation screening. While not a primary screen hit, hsa-miR-34a-5p was included for further validation as described in the text. A functional secondary screen found that 51/73 hits strongly inhibited nucleolar rRNA biogenesis via 5-EU incorporation. Additional mechanistic assays for pre-rRNA transcription or processing, global translation, and nucleolar stress were carried out for a subset of 15 rigorously-selected hits. Further mechanistic studies were performed on two MIR-28 family siblings, hsa-miR-28-5p and hsa-miR-708-5p. (B) Representative images from the screen showing 3 top hits alongside the negative control (non-targeting, siNT) and positive controls for the one-nucleolus (siNOL11) or 5+ nucleoli (siKIF11) phenotypes. hsa-miR-548-an and hsa-miR-708-5p are hits with the one-nucleolus phenotype while hsa-miR-629-3p is a hit with the 5+ nucleoli phenotype. The DNA stain (Hoechst) is shown in blue and fibrillarin (FBL) antibody detection is shown in red. Scale bars, 10 μm. (**C**). Double flashlight plot for one-nucleolus hit selection. A total of 64 one-nucleolus hits were called using cutoffs for the top 2.5% of mean one-nucleolus percent effect and at least fairly strong hits where SSMD ≥ 1.645 as described in the text. Hits in MirGeneDB are shown in blue, while hits not in MirGeneDB are shown in red. Data were graphed in GraphPad Prism 8. (**D**) Double flashlight plot for 5+ nucleoli hit selection. A total of nine 5+ nucleoli hits were called using cutoffs for the top 1.0% of mean 5+ nucleoli percent effect and the same SSMD cutoff as above. Hits in MirGeneDB are shown in blue, while hits not in MirGeneDB are shown in red. Data were graphed in GraphPad Prism 8.

We took several steps to ensure our screen’s reproducibility and minimize false positives. We conducted the primary screen in biological triplicate, enabling robust calculation of one-nucleolus percent effect and 5+ nucleoli percent effect for each microRNA mimic. To assist with hit selection, we also calculated the strictly standardized mean difference (SSMD) for each microRNA mimic.

SSMD is a more robust estimator of hit effect size than percent effect alone, as it also incorporates information on sample size and reproducibility (variance) while simultaneously controlling the false positive and false negative rates (92). Using these two cutoffs in concert allowed us to pick the strongest, most replicable hits. The percent effect cutoff was set at the top 2.5% for the one-nucleolus effect and at the top 1.0% for the 5+ nucleoli effect; the SSMD cutoff was set at 1.645 for both phenotypes, which allows identification of hits that are at least fairly strong (58). The median signal-to-background (S/B) for the controls was 2.74 or 2.64 for the one-nucleolus screen or the 5+ nucleoli screen, respectively. The median Z’ factor for the one-nucleolus screen was 0.41, although the median Z’ factor for the 5+ nucleoli screen was unfavorable at -0.43. Despite poor separation of controls for the 5+ nucleoli screen, reflected in the low median Z’ factor, we felt confident that imposing a stricter percent effect cutoff and maintaining the strong SSMD cutoff would enable us to identify hits with reproducible increases in nucleolar number.

Using stringent cutoffs for mean percent effect and SSMD, the nucleolar number primary screen identified 64 one-nucleolus hits and 9 5+ nucleoli hits (**Figure 1C-D, Supplementary Tables 2-3**). Of these hits, we found that 24/64 one-nucleolus hits and 2/9 5+ nucleoli hits were present in the MirGeneDB database, which catalogs evolutionarily-conserved microRNAs (**Figure 1C-D**) (52). The SSMD cutoff approach allowed us to ignore several less reproducible hits with otherwise high mean percent effect values, mostly from the 5+ nucleoli side of the screen (**Figure 1C-D**, bottom right quadrant of each graph). This total of 73 hits equates to an overall 2.8% hit rate. Inspection of images from top hits including hsa-miR-548an, hsa-miR-708-5p, and hsa-miR-629-3p (**Figure 1B**) confirmed that the relevant phenotype for each hit was observed. We performed a validation screen with replicates for the 73 hits, where each hit was picked from the original microRNA mimic library and rescreened on a new plate for a change in nucleolar number. Overall, 71/73 hits (97%) passed validation (**Supplementary Figure 1, Supplementary Table 3**). Interestingly, our hits show a statistically-significant difference in chromosomal distribution than the natural distribution of 1918 microRNA genes in the human genome cataloged by miRBase (Chi-squared test *p* < 0.031; **Supplementary Tables 4-5**), which may warrant additional future investigation. The standardized residuals indicate the 71 hits are most enriched for loci on chromosomes 20 and 9 (**Supplementary Table 5**).

### Novel microRNA negative regulators of ribosome biogenesis preferentially target transcripts encoding proteins in the nucleolus or involved in cell cycle regulation

We hypothesized that the microRNA mimic hits from the primary screen would be more likely to target genes whose product is nucleolar or is involved in processes affecting the nucleolus. We sought to combine bioinformatic databases containing microRNA:target RNA pairs, MCF10A RNA expression data, and catalogs of known nucleolar proteins to test this hypothesis. To investigate the hits’ targets, we utilized TarBase 8, a catalog of over 670,000 experimentally-validated microRNA:target RNA interactions collected from the literature (66). We filtered TarBase 8 to only include 373,890 human microRNA:target interactions with a “down” regulatory relationship (**Figure 2A**). We assembled an MCF10A RNA expression dataset from 4 independent RNAseq experiments (BioProject accessions PRJNA290557, PRJNA384982, PRJNA530983, PRJNA647393) to determine which RNAs were expressed in this cell line. We discovered 20,345 genes bearing one or more transcripts with a normalized (zTPM) expression value greater than -3 in at least one RNAseq experiment (**Supplementary Table 8**), supporting expression for each of these genes (59). We used our RNA expression dataset to further filter out all target genes that were not expressed in MCF10A cells from TarBase 8, after which 1,074 microRNAs and 351,983 microRNA:target interactions remained (**Figure 2A**). Furthermore, we merged three nucleolar protein databases (63–65) to create a nucleolar protein reference metadatabase containing 3,490 unique nucleolar proteins (**Figure 2A, Supplementary Table 9**). Using these three datasets, we labeled all microRNA:target interactions potentially present in MCF10A cells that contain targets encoding nucleolar proteins. Only 32/71 (45.1%) of the microRNA hits had at least one experimentally-validated target in TarBase 8 (**Figure 2A**); however, this figure is consistent with the fact that only 1,074/2,603 (41.3%) microRNAs in the primary screening library are represented in TarBase 8.

**Figure 2.**
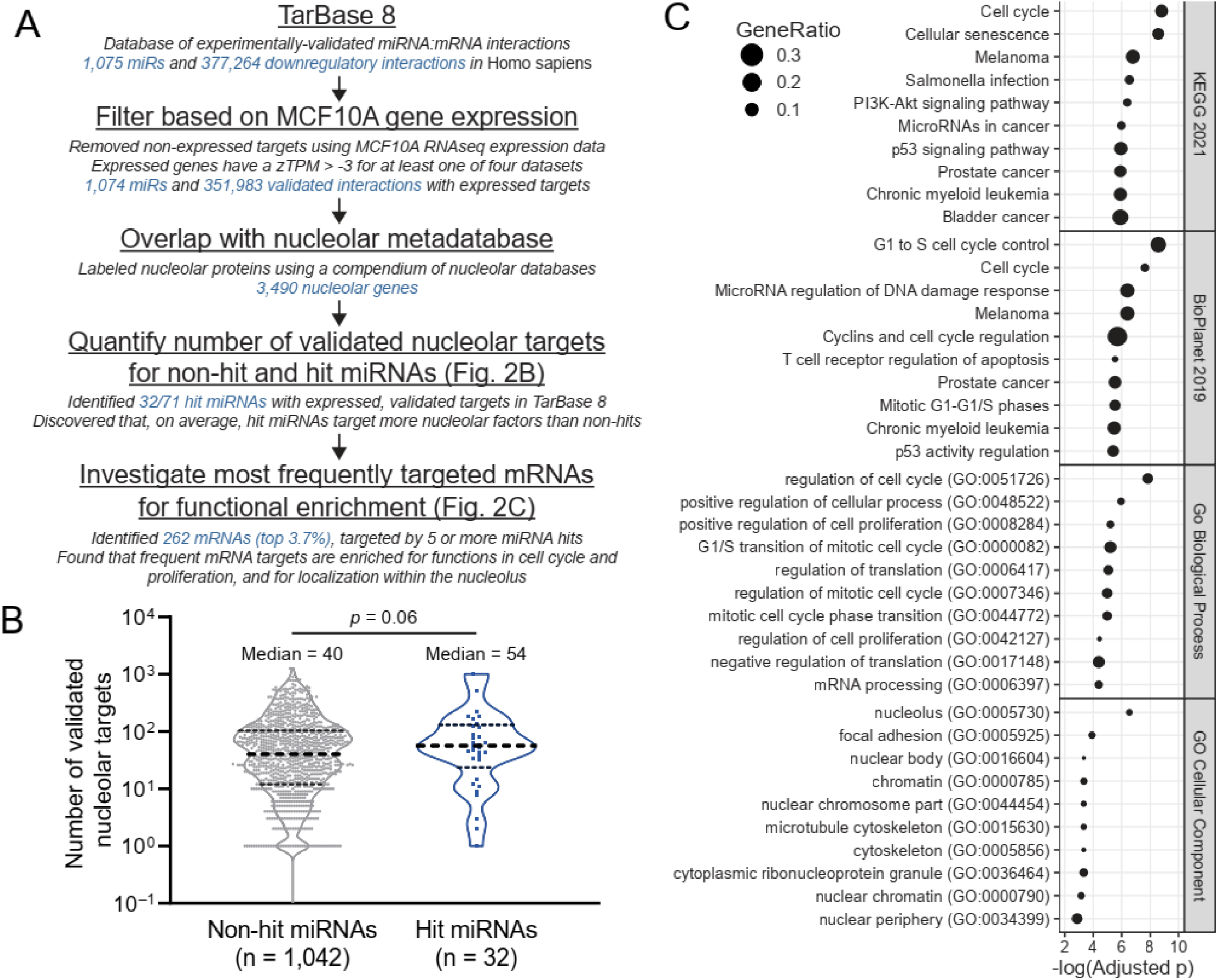
The novel microRNA hits preferentially target genes encoding proteins with nucleolar localization or with functions in cell cycle control. (**A**) Bioinformatics workflow. TarBase 8 contained 520,410 validated microRNA:RNA target interactions across 1,075 mature human microRNAs. TarBase 8 was filtered for genes expressed in MCF10A cells, using a cutoff of more than -3 zTPM (normalized TPM), leaving 351,983 validated interactions. Targets were labeled for nucleolar localization based on our nucleolar proteome metadatabase containing 3,490 nucleolar proteins. MicroRNAs were grouped based on primary screen hit status, with 32/71 primary screen hits having one or more validated, expressed targets in MCF10A cells. The number of validated, expressed, nucleolar targets was calculated for hit and non- hit microRNAs. Conversely, the number of hit microRNAs targeting each gene was calculated, and all genes targeted by 5 or more hits (262, top 3.7%) were analyzed for enrichment. (**B**) Log_10_-scale plot indicating the number of validated nucleolar targets expressed in MCF10A cells for hit and non-hit microRNAs in TarBase 8. The median number of nucleolar targets per hit microRNA is 54, which is greater than the non-hit median of 40. Data were graphed and analyzed with an unpaired two-sided t-test in GraphPad Prism 8. (**C**). Enrichment plots for 262 genes targeted by 5 or more of the microRNA hits. Plots indicate −log_10_(adjusted *p*) on the x-axis and the gene ratio as the marker size. Enrichment analysis was conducted with Enrichr, and plots were made in R. Enrichment databases: Kyoto Encyclopedia of Genes and Genomes (KEGG) 2021; NCATS BioPlanet of Pathways 2019; Gene Ontology (GO) Biological Process 2018; GO Cellular Component 2018.

To investigate our hypothesis that hit microRNAs preferentially target transcripts encoding nucleolar proteins, we compared the 32 hit microRNAs to the remaining 1,042 non-hit microRNAs in our filtered, annotated TarBase 8 database (**Figure 2B**). We counted the number of transcripts coding nucleolar proteins that are targeted by each microRNA. We find that the median number of nucleolar targets for the hit microRNAs is 54, compared to the median value of 40 for 1,042 non-hit microRNAs, supporting our hypothesis (**Figure 2B**). We also identified 262 genes expressed in MCF10As that are targeted by at least 5 of the novel microRNA hits, representing the top 3.7% of genes most frequently targeted by the hits (**Figure 2A, Supplementary Table 10**). GO analysis of these genes revealed enrichment for encoded functions in cell cycle regulation, TP53 signaling, cellular proliferation, and for localization within the nucleolus (**Figure 2C**). We also identified 135 genes expressed in MCF10As that are targeted by at least 5 of the 24 novel microRNA hits present in TarBase 8 and MirGeneDB (52); these genes were similarly enriched for nucleolar localization and regulators of the cell cycle and growth (**Supplementary Figure 2, Supplementary Table 10**).

Altogether, these data support the hypothesis that the novel microRNA hits preferentially target transcripts encoding proteins localized to the nucleolus or involved in cell cycle regulation.

### A majority of novel microRNA negative regulators of ribosome biogenesis strongly inhibit nucleolar rRNA biogenesis

To more directly determine the extent to which the novel microRNA hits can abrogate nucleolar function, we harnessed our laboratory’s high-throughput assay for nucleolar rRNA biogenesis inhibition (55). Briefly, we previously optimized and miniaturized a nascent rRNA assay using 5- ethynyl uridine (5-EU) metabolic labeling to quantify changes in nucleolar 5-EU signal by co-staining for the nucleolar protein FBL/fibrillarin (**Figure 3A**). Our assay is sensitive to defects in pre-rRNA transcription, processing, or modification, which we collectively refer to as nucleolar rRNA biogenesis (55). We established an empirical cutoff for nucleolar rRNA biogenesis inhibition of 50%, as we discovered that depletion of almost all RB factors we tested during validation caused inhibition at or above this value (55). Furthermore, factors involved in both pre-rRNA transcription and processing typically caused the highest percent inhibition values, followed by factors only involved in pre-rRNA transcription, pre-rRNA processing, or pre-rRNA modification, respectively.

**Figure 3.**
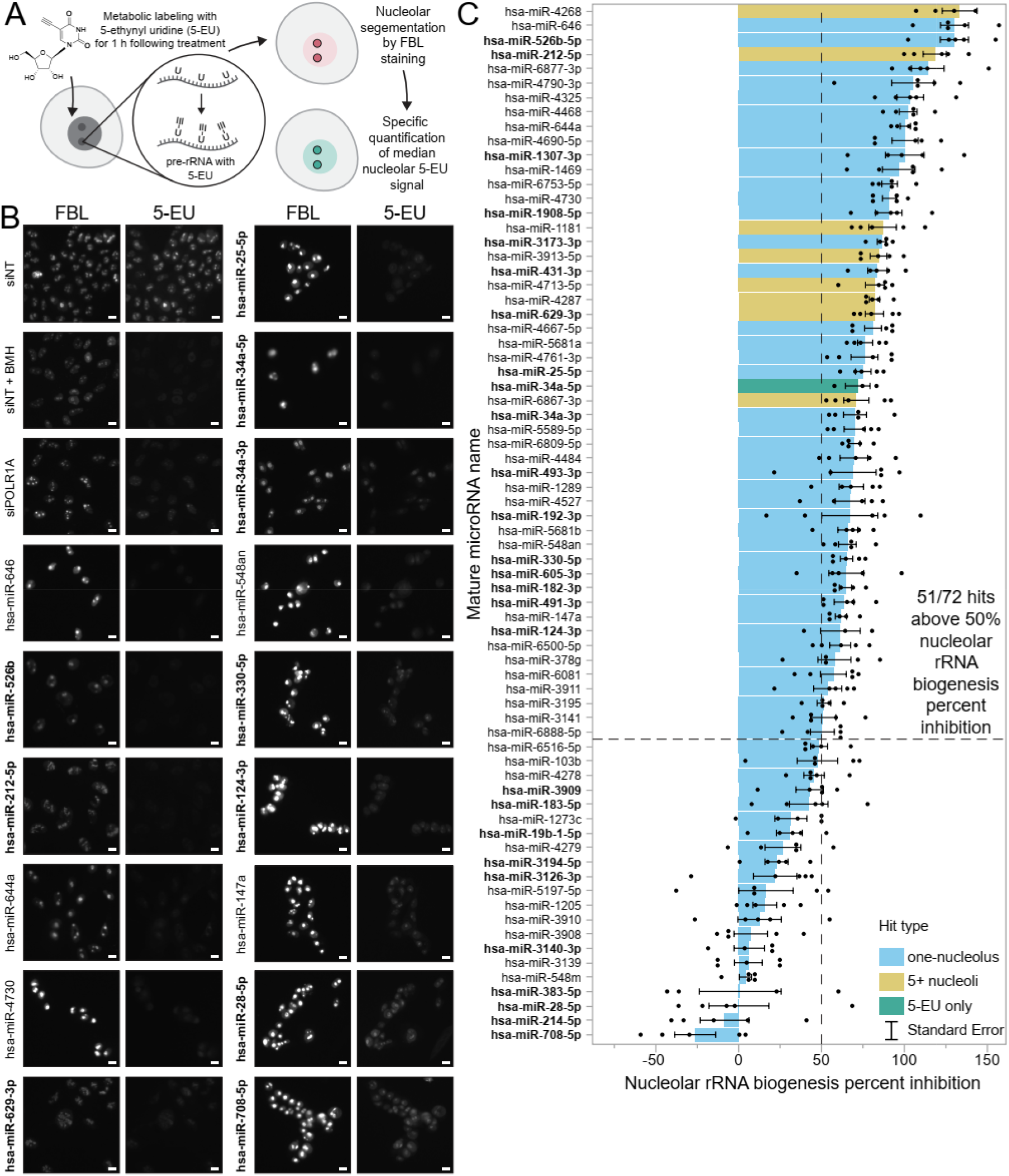
A secondary screen reveals 51/72 hits strongly inhibit nucleolar rRNA biogenesis. (**A**). Schematic for the nucleolar rRNA biogenesis inhibition assay (55). Following 72 h of RNAi transfection, MCF10A cells were labeled for 1 h with 1 mM 5-ethynyl uridine (5-EU). Cells were fixed and immunostained for the nucleolar protein FBL and for 5-EU, then imaged. CellProfiler was used to segment nucleoli and calculate the median 5-EU intensity for all nucleoli per well, enabling calculation of the nucleolar rRNA biogenesis percent inhibition. (**B**) Representative images of control- and hit-treated MCF10A cells following 5-EU incorporation. FBL immunostaining and 5-EU click labeling are shown as separate channels. Scale bars, 10 μm. siNT is the non-targeting negative control siRNA. siNT + BMH is siNT-transfected cells treated with 1 μM BMH-21 for 1 h before and during 5-EU incorporation. siPOLR1A is the POLR1A (RPA194) knockdown positive control. (**C**). Nucleolar rRNA biogenesis percent inhibition values for 72 microRNA mimic hits. A total of 51/72 hits caused at least 50% inhibition of nucleolar rRNA biogenesis, surpassing the assay’s empirical cutoff (55). siNT negative control is set to 0% inhibition, and siPOLR1A positive control is set to 100% inhibition. Hit names are bolded for MirGeneDB members, and bar graphs are colored according to their primary screen phenotype; hsa-miR-34a-5p was not a primary screen hit but was included as described in the text. Mean ± SEM are shown alongside individual data points (*n* = 5 biological replicates). Data were graphed in JMP.

We conducted a secondary screen with five biological replicates for nucleolar rRNA biogenesis inhibition on all 71 microRNA mimics that passed primary screen validation as well as one additional microRNA mimic in the original library as a positive control, hsa-miR-34a-5p. We chose to include both hsa-miR-34a strands, as the *MIR34A* locus has been implicated in Wnt-mediated control of RB (93), and hsa-miR-34a-5p targets the *RMRP* RNA which is critical for pre-rRNA processing (94,95).

Following treatment, cells were fixed and stained for FBL and 5-EU incorporation (**Figure 3B**), before image processing and quantification were completed with CellProfiler. As expected, MCF10A cells treated with siNT had robustly active nucleoli (**Figure 3B**, siNT panel), while positive control cells depleted of the RNAP1 subunit POLR1A (siPOLR1A) showed strongly decreased nucleolar rRNA biogenesis (**Figure 3B**, siPOLR1A panel) (55). Acute treatment with BMH-21, a known small molecule inhibitor of RNAP1 activity (96,97), at 1 μM for 1 h also eliminated nucleolar 5-EU signal as expected (55) (**Figure 3B**, siNT + BMH panel). For the secondary screen replicates (*n* = 5), the median Z’ factor was 0.27 and the median S/B was 1.88, indicating acceptable separation of controls. In additional experiments (*n* = 2), we assayed the ability of the 27 evolutionarily-conserved MirGeneDB hits to inhibit nucleolar rRNA biogenesis in two other human cell lines, hTERT RPE-1 and HeLa (**Supplementary Figure 3**).

Remarkably, the full secondary screen in MCF10A cells indicated that 51/72 (70.8%) microRNA mimic hits assayed caused at least a 50% inhibition of nucleolar rRNA biogenesis (**Figure 3C, Supplementary Table 3**). Notably, all eight 5+ hits tested strongly inhibited nucleolar rRNA biogenesis, with a mean percent inhibition of 92.6%. Comparing our results in MCF10A cells to those in hTERT RPE-1 and HeLa cells, we found that 17/27 (63.0%) MirGeneDB hits tested caused at least a 50% inhibition of nucleolar rRNA biogenesis in two or more of these cell lines (**Supplementary Figure 3**). These data support the hypothesis that most microRNA hits from the primary screen significantly disrupt human nucleolar rRNA biogenesis, the main function of the nucleolus, in multiple human cell lines.

### A diverse subset of 15 microRNA hits was chosen for mechanistic follow-up

We chose a tractable subset of 15 microRNA hits to further study for their specific effects on crucial RB subprocesses, including pre-rRNA transcription, pre-rRNA processing, and global protein synthesis. We prioritized selecting a diverse group of microRNAs that were representative of differences in key variables observed in the screening campaign, but also were considered to be authentic, valid microRNAs by sequencing and evolutionary analyses. To this end, we conducted principal component analysis (PCA) to visualize screening and bioinformatic data in a dimension-reduced format (**Figure 4A-B**). A major cluster appeared containing most one-nucleolus hits with relatively few validated nucleolar targets, accompanied by outliers that either were 5+ nucleoli hits or had a high number of validated nucleolar targets (**Figure 4A**). To minimize the potential for studying biological false positives, we also classified hits according to their evolutionary conservation, as cataloged by MirGeneDB (52), or their sequencing read quality consistency across 28,866 small RNAseq experiments (98). The latter analysis accounted for the compliance of small RNAseq reads with Dicer processing rules, particularly the minimal variability of microRNA sequences at the 5’ terminus, and for coverage across the microRNA precursor transcriptome annotation.

**Figure 4.**
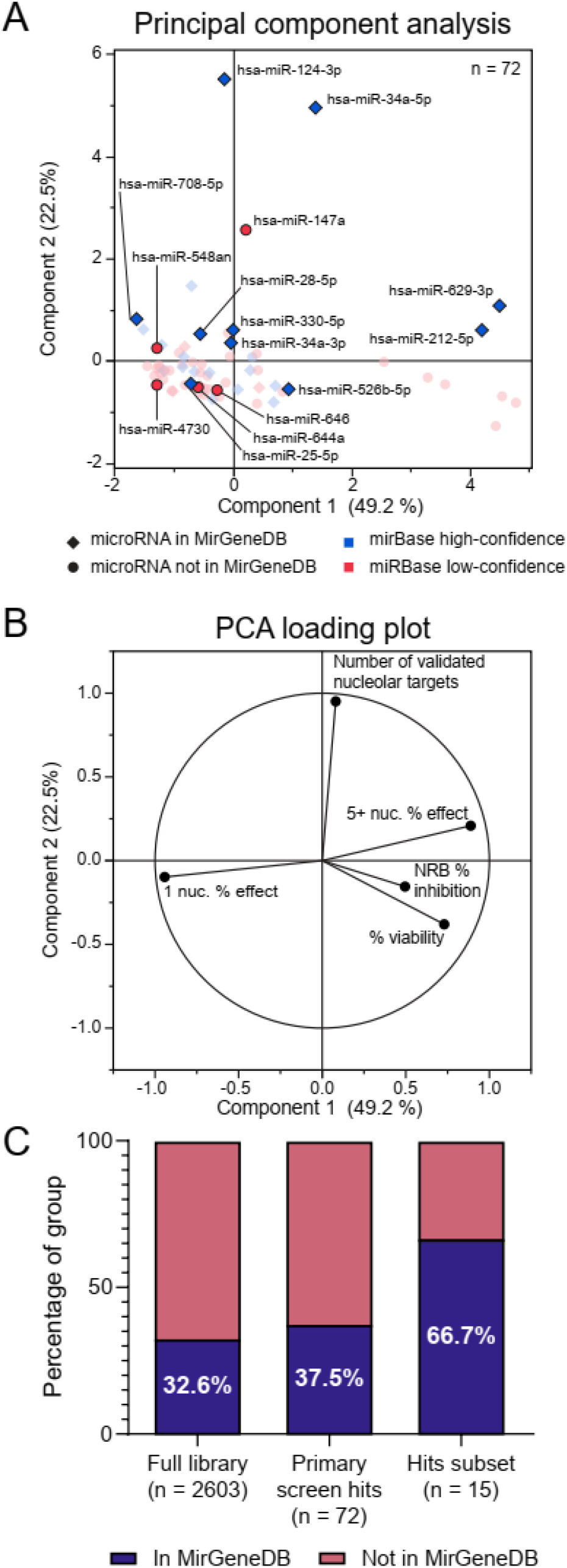
A subset of 15 microRNA hits were rigorously selected for additional mechanistic validation. (A) Principal component analysis (PCA) of 72 hits. The 15 selected subset hits are indicated with their mature microRNA name and are highlighted on the plot. Membership in MirGeneDB (52) or classification as high-confidence or low-confidence miRBase (98) is labeled with each hit’s marker shape or color, respectively. Five variables were used for PCA including one-nucleolus and 5+ nucleoli percent effect, percent viability, nucleolar rRNA biogenesis (NRB) percent inhibition, and number of validated nucleolar targets for each hit in TarBase 8. Percentages in axis labels denote the proportion of variance explained by each PCA Component. (B) Loading plot describing contribution of 5 quantitative variables to PCA Components 1 and 2 from above. Percentages in axis labels denote the proportion of variance explained by each PCA Component. (C) Stacked bar graphs of the percent of each listed group of microRNA mimics present in MirGeneDB.

For further mechanistic assay validation, we combined our PCA analysis, microRNA conservation data, and literature curation to manually select 12 one-nucleolus hits, two 5+ nucleoli hits, and hsa-miR-34a-5p, which was not a hit in the primary screen but did significantly inhibit nucleolar rRNA biogenesis (**Supplementary Tables 6-7**). The median nucleolar rRNA biogenesis percent inhibition of these hits was 72.2% with a range of -26.3% to 130.3%. Of these hits, 10 were recorded in MirGeneDB and classified as “High Confidence” microRNAs annotated in miRBase (98), while 5 were not present in MirGeneDB and were classified as “Low Confidence” microRNAs in miRBase (98) (**Supplementary Tables 3, 6, and 7**). Therefore, our curated hit selection process significantly enriched for conserved microRNAs present in MirGeneDB, as compared to the full microRNA mimic library and set of primary screen hits (**Figure 4C**). Additionally, 14/15 hits passed a tripartite filter for sequencing read quality consistency (98). Two microRNA hits from the MirGeneDB MIR-28 family (52), hsa-miR-28-5p and hsa-miR-708-5p, were included in the subset.

### A subset of microRNA hits dysregulates pre-rRNA transcript levels and rDNA promoter activity

Since many microRNA hits caused strong inhibition of nucleolar rRNA biogenesis in the secondary 5-EU screen, we hypothesized that these hits might dysregulate pre-rRNA transcription directly by targeting the 47S primary transcript. We tested this hypothesis by comparing the number of predicted canonical seed binding sites on the 47S pre-rRNA transcript (RefSeq transcript NR_046235.3) for each of the 72 hits with its corresponding mean nucleolar rRNA biogenesis percent inhibition (**Supplementary Figure 4A; Supplementary Table 13**). Surprisingly, we did not observe a strong correlation, with several hits having a high nucleolar rRNA biogenesis percent inhibition and zero predicted 47S binding sites.

To experimentally test the 15 subset hits’ effects on RNAP1 transcription, we measured the steady-state levels of the 45S pre-rRNA transcript as a proxy for transcription after treatment with the subset of microRNA mimics or control siRNAs by RT-qPCR. (**Supplementary Figure 4B**). While depletion of the positive control transcription (t)UTP NOL11 led to a stark decrease in 45S levels versus siNT as expected (91), none of the microRNA hits statistically-significantly altered 45S levels (**Supplementary Figure 4C**). However, 4 microRNA hits (hsa-miR-34a-5p, hsa-miR-212-5p, hsa-miR-330-5p, hsa-miR-526b-5p) caused a mean decrease in 45S levels of at least 50% [less than -1 log_2_(45S transcript levels)]. However, considerable experimental noise with this assay may mask the true effect of these microRNA mimics on altering 45S transcript levels.

To further interrogate pre-rRNA transcription, we conducted a dual-luciferase reporter assay for RNAP1 promoter activity (78,99) after microRNA mimic expression. The assay uses a firefly luciferase reporter under the control of the rDNA promoter and a *Renilla* luciferase reporter constitutively driven by a CMV promoter to normalize for transfection efficiency (**Supplementary Figure 4D**). Strikingly, we found that treatment with the 15 microRNA hits had diverse effects on RNAP1 promoter activity: 3 microRNA mimics (hsa-miR-147a, hsa-miR-526b, hsa-miR-548an) caused a decrease in RNAP1 promoter activity; 5 microRNA mimics (hsa-miR-34a-5p, hsa-miR-124- 3p, hsa-miR-330-5p, hsa-miR-629-3p, hsa-miR-646) caused an increase in RNAP1 promoter activity; and the other 7 microRNA mimics did not cause a significant effect (**Supplementary Figure 4E**).

Compared to mock and siNT negative controls, siNT treatment followed by positive control 1 μM BMH-21 dosage significantly decreased RNAP1 promoter activity, while positive control NOL11 depletion caused a modest but statistically-insignificant defect (**Supplementary Figure 4E**).

Conversely, depletion of the cytoplasmic 60S ribosomal subunit protein RPL4 had no effect on RNAP1 promoter activity (**Supplementary Figure 4E**). These results indicate that the microRNA hits do not reliably affect pre-rRNA transcription as measured by 5-EU incorporation or 45S transcript levels, while they may upregulate, downregulate, or have no effect on RNAP1 promoter activity.

Future experiments may be required to better understand the interplay between microRNA activity and pre-rRNA transcription.

### A subset of microRNA hits dysregulates maturation of the 30S pre-rRNA precursor

Given the inconclusive ability of the microRNA hit subset to consistently regulate pre-rRNA transcription, we also hypothesized that these hits could affect pre-rRNA processing, another step in nucleolar rRNA biogenesis (55). We carried out northern blotting for all 15 microRNA mimic hits in the subset, probing for pre-rRNA processing intermediate molecules containing probes for either internal transcribed spacer 1 (ITS1, P3 probe for pre-40S defects) or ITS2 (P4 probe for pre-60S defects) (**Figure 5A**). Surprisingly, ITS1 blots broadly demonstrated that most microRNA hits dysregulated maturation of the 30S pre-rRNA precursor (**Figure 5B-C**). Ratio Analysis of Multiple Precursors calculations [RAMP, (77)] indicated 3 major clusters of pre-rRNA processing defects caused by the microRNA hits, namely, no change (n.c.), a 30S down cluster, and a 30S up/21S down cluster (**Figure 5C-D**). The last cluster contained two subclusters, one with hits causing a moderate 30S processing defect (“30S↑, 21S↓” in **Figure 5D**) and another with hsa-miR-28-5p and hsa-miR-708-5p which caused a severe 30S processing defect (“30S↑, 21S↓↓” in **Figure 5D**). We also examined the extent to which ITS2 processing was dysregulated by the subset of microRNA hits (**Supplementary Figure 5A**). We discovered that hsa-miR-708-5p and the two 5+ nucleoli hits, hsa-miR-212-5p and hsa-miR-629-3p, each caused a mild increase in 32S levels and a mild decrease in 12S levels; additionally, hsa-miR-330-5p mildly attenuated levels of the 32S and 12S precursors (**Supplementary Figure 5B-C**). The remaining microRNA hits did not cause a significant change in ITS2-containing precursor levels.

**Figure 5.**
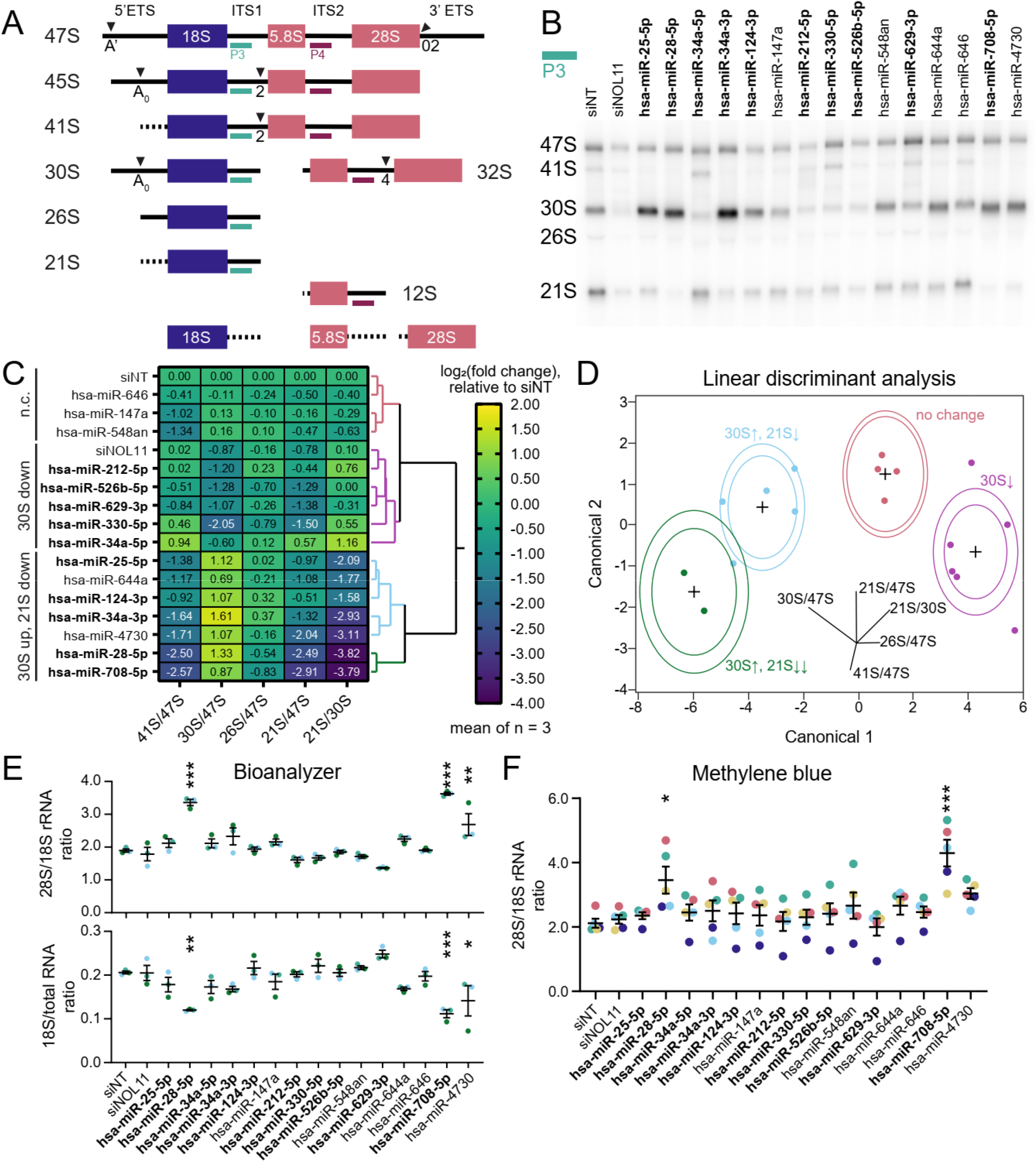
Most subset microRNA hits dysregulate pre-18S pre-rRNA processing. **(A)** Simplified diagram of human pre-rRNA processing intermediates. Mature 18S, 5.8S, and 28S rRNA regions are shown as blue (pre-40S) or red (pre-60S) rectangles. Intermediate names are indicated on the left, and external and internal transcribed spacers (ETS or ITS, solid black lines) are labeled at the top. Cleavage sites are labeled with their name and represented with triangles. Dotted lines signify transcribed spacer regions digested by exonucleases. Northern blot probes P3 (teal, ITS1) or P4 (dark red, ITS2) are shown at each pre-rRNA intermediate that they bind. (B) Representative ITS1 (probe P3) northern blot of 3 μg of total RNA isolated from control- or hit- treated (as indicated) MCF10A cells. Pre-rRNA processing intermediates are labeled on the left. Images were quantified using Bio-Rad Image Lab. siNT is the non-targeting negative control and siNOL11 is the positive control. Hit names are bolded for MirGeneDB members. (C) Clustered heatmap showing log_2_-transformed Ratio Analysis of Multiple Precursor [RAMP (77)] calculations for microRNA hits, normalized to siNT negative control. Values represent mean log_2_- scale RAMP ratio for *n* = 3. Clusters: no change (n.c., red); 30S down (magenta); 30S up, 21S down (mild defect, blue and severe defect, green). RAMP ratios were calculated in Microsoft Excel. Four clusters were assigned using hierarchical Ward clustering in JMP, and data were graphed in GraphPad Prism 8. Hit names are bolded for MirGeneDB members. (D) Linear discriminant analysis (LDA) of four 30S pre-rRNA processing defect clusters from (**C**) above. Cluster colors are the same as in (**C**). Canonical component biplot rays are shown in the graph. Ellipses represent 50% (outer) and 95% (inner) confidence levels. Hit names are bolded for MirGeneDB members. Data were graphed in JMP. (E) Bioanalyzer analysis for 1 μg of total RNA isolated from control- or hit-treated MCF10A cells. Top, the 28S/18S mature rRNA ratio; bottom, 18S mature rRNA/total RNA ratio. Mean ± SEM are shown alongside individual data points, colored by replicate. Hit names are bolded for MirGeneDB members. Data were graphed and analyzed by ordinary one-way ANOVA with multiple comparisons against siNT (non-targeting negative control) and Holm-Šídák correction in GraphPad Prism 8. **\***, *p* < 0.05; **\*\***, *p* < 0.01; **\*\*\***, *p* < 0.001. (F) Methylene blue (MB) analysis of the 28S/18S mature rRNA ratio. Northern blots from (**B-C**) were stained with MB, imaged, and quantified using Bio-Rad Image Lab. Mean ± SEM are shown alongside individual data points, colored by replicate. Hit names are bolded for MirGeneDB members. Data were graphed and analyzed by ordinary one-way ANOVA with multiple comparisons against siNT (non-targeting negative control) and Holm-Šídák correction in GraphPad Prism 8. **\***, *p* < 0.05; **\*\***, *p* < 0.01; **\*\*\***, *p* < 0.001.

We conducted Bioanalyzer electrophoresis to define the ability of the microRNA hits to modulate steady-state levels of mature 28S and 18S rRNAs (**Figure 5A**). Bioanalyzer quantification revealed an increase in the mature 28S/18S rRNA ratios for 3 microRNA hits, hsa-miR-28-5p, hsa-miR-708-5p, and hsa-miR-4730 (**Figure 5E**), portending a defect in 18S maturation. Indeed, calculation of the ratio of mature 18S rRNA to total RNA from the electropherogram data showed stark decreases in 18S levels following treatment with any of these three microRNA mimics (**Figure 5E**). Methylene blue staining corroborated the increase in 28S/18S ratio for hsa-miR-28-5p and hsa-miR-708-5p (**Figure 5F**), although this technique has lower precision than Bioanalyzer quantification.

All three microRNA mimics with deficient mature 18S rRNA levels were in the severe 30S processing defect cluster, consistent with the pre-rRNA processing pathway (**Figure 5A**), and hsa-miR-4730 clustered at the interface between the moderate and severe 30S defect subclusters (**Figure 5D**). The other 12 microRNA mimic hits did not cause a significant change in mature rRNA levels. Together these results reveal, for the first time, the dysregulatory potential of microRNAs to interfere with major steps in pre-rRNA processing.

### A subset of microRNA hits decreases global translation

We also hypothesized that overexpression of the microRNA hits might repress global translation. We used the SUnSET puromycin labeling assay (80) to quantify the extent to which each microRNA hit could inhibit global protein synthesis. Following a 72 h transfection period, MCF10A cells were metabolically labeled for 1 h with 1 μM puromycin, which is incorporated into nascent peptides. Total puromycin was measured as a proxy for global translation rate by immunoblotting total protein with an α-puromycin primary antibody. Treatment with 14/15 (93.3%) of the microRNA hits significantly decreased global protein synthesis relative to the negative non-targeting siRNA control (**Figure 6A-B**). A similar decrease was observed for the positive control, an siRNA depleting the 60S ribosomal protein RPL4 (**Figure 6A-B**). In total, these results indicate that nearly all of the subset microRNA hits inhibit global protein synthesis, the ultimate endpoint of ribosome biogenesis.

**Figure 6.**
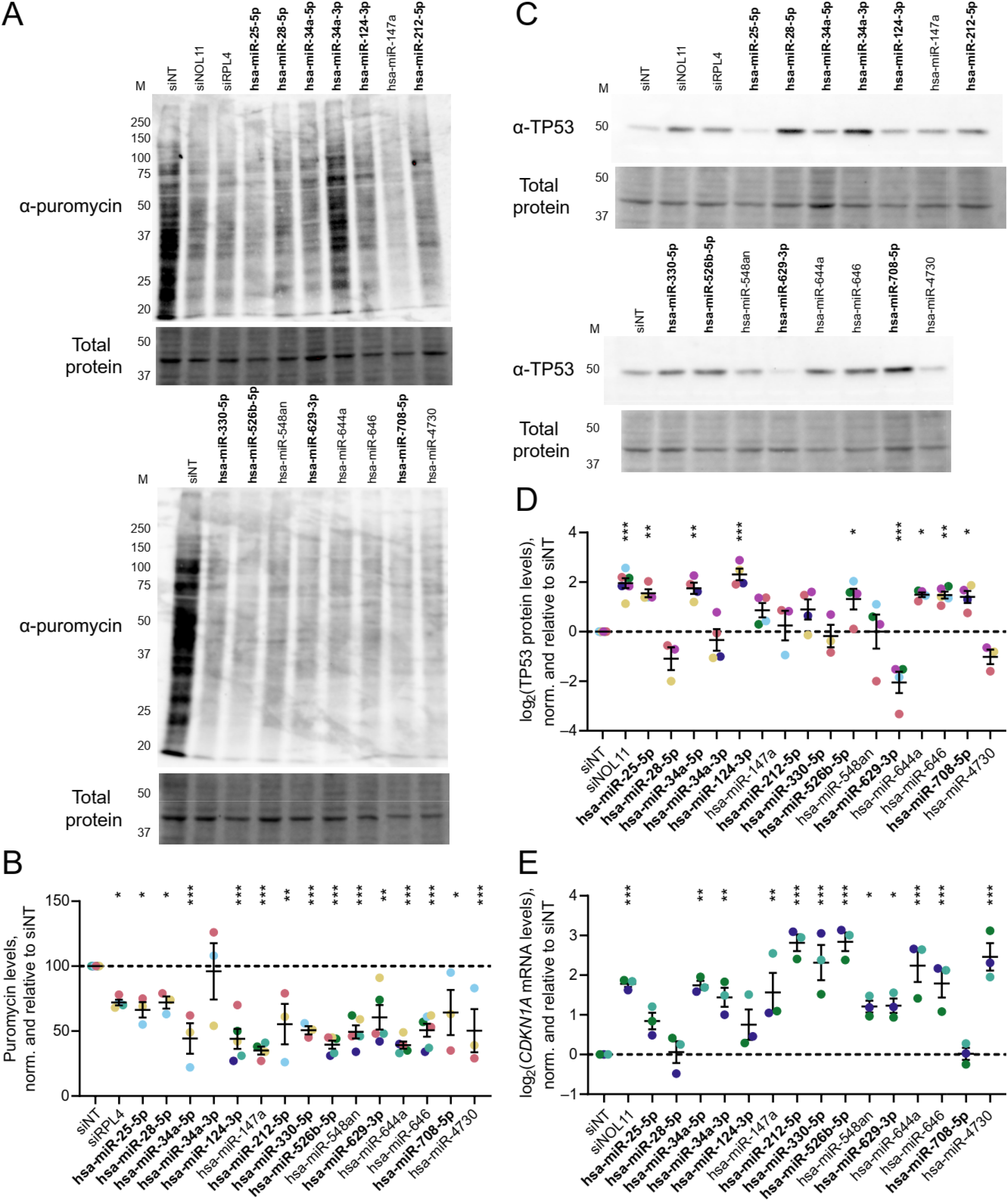
MicroRNA hits inhibit global protein synthesis and dysregulate levels of the cell cycle regulators TP53 and *CDKN1A*. (A). Representative examples of the SUnSET puromycin incorporation assay (53,80) immunoblots of total protein isolated from control- or hit-treated MCF10A cells. α-puromycin is an immunoblot for puromycin incorporation as a proxy for global protein synthesis. Total protein is the trichloroethanol total protein stain loading control. M, molecular weight marker lane in kDa. The images were quantified with Bio-Rad Image Lab. Hit names are bolded for MirGeneDB members. (B). Quantification of global protein synthesis levels from (**A**) above. Mean ± SEM are shown alongside individual data points, colored by replicate (at least 3 replicates per condition, with all replicates shown in the graph). Hit names are bolded for MirGeneDB members. The data were normalized to siNT, then graphed and analyzed by ordinary one-way ANOVA with multiple comparisons against siNT and Holm-Šídák correction in GraphPad Prism 8. **\***, *p* < 0.05; **\*\***, *p* < 0.01; **\*\*\***, *p* < 0.001. (C). Representative examples of TP53 immunoblots of total protein isolated from control- or hit- treated MCF10A cells. α-TP53 is the blot for TP53. Total protein is the trichloroethanol total protein stain loading control. M, molecular weight marker lane in kDa. Hit names are bolded for MirGeneDB members. (D). Log_2_-scale quantification of TP53 protein levels from (**C**) above. Mean ± SEM are shown alongside individual data points, colored by replicate (at least 3 replicates per condition, with all replicates shown in the graph). Hit names are bolded for MirGeneDB members. Data were normalized to siNT, then graphed and analyzed by ordinary one-way ANOVA with multiple comparisons against siNT and Holm-Šídák correction in GraphPad Prism 8. **\***, *p* < 0.05; **\*\***, *p* < 0.01; **\*\*\***, *p* < 0.001. (E). Log_2_-scale RT-qPCR analysis of *CDKN1A* (*p21*) mRNA levels from control- or hit-treated MCF10A cells. The data from 3 biological replicates were normalized to the 7SL RNA abundance as a loading control, then to siNT for comparison using the ΔΔC_T_ method. Mean ± SEM are shown alongside individual data points, colored by replicate. Hit names are bolded for MirGeneDB members. Data were analyzed by ordinary one-way ANOVA with multiple comparisons against siNT and Holm- Šídák correction in GraphPad Prism 8. **\***, *p* < 0.05; **\*\***, *p* < 0.01; **\*\*\***, *p* < 0.001.

### A subset of microRNA hits alters levels of TP53 or *CDKN1A* (*p21*)

Since treatment with many microRNA mimic hits significantly inhibited nucleolar rRNA biogenesis and normal pre-rRNA processing, we hypothesized that treatment with the hits may activate the nucleolar stress response via TP53 stabilization and *CDKN1A* (*p21*) upregulation. The nucleolar stress response is induced following disruption of RB subprocesses or the normal tripartite nucleolar structure (17,19,100). Mechanistically, 5S RNP proteins including the 60S ribosomal proteins RPL5 or RPL11 can bind and sequester MDM2, the E3 ubiquitin ligase targeting TP53 for constitutive degradation. De-repression of TP53 then upregulates *CDKN1A* (*p21*) expression, which acts in concert with TP53 to arrest the cell cycle and initiate apoptosis. By harnessing the WT TP53 status of MCF10A cells, we tested the first part of our hypothesis by immunoblotting for steady-state TP53 levels to assess how microRNA hit overexpression affected nucleolar stress response induction (**Figure 6C**). Six of the 15 microRNA hits stabilized TP53 levels, while surprisingly, hsa-miR-629-3p significantly decreased steady-state levels of TP53 (**Figure 6C-D**). The remaining 8/15 microRNAs did not cause significant dysregulation of TP53 levels. Depletion of either of the positive controls, tUTP factor NOL11 or the 60S ribosomal protein RPL4, strongly stabilized TP53 as expected (**Figure 6C-D**).

We also investigated how the microRNA hits affected *CDKN1A* (p21) mRNA transcript levels by RT-qPCR. Again, depletion of the positive control NOL11 robustly increased *CDKN1A* levels as compared to siNT, while 11/15 microRNA mimics also upregulated *CDKN1A* to a statistically-significant degree (**Figure 6E**). The TP53 and *CDKN1A* data largely concur, although there are two notable discrepancies. First, hsa-miR-28-5p and hsa-miR-708-5p caused strong TP53 stabilization yet did not elicit measurable *CDKN1A* upregulation, which we address in the next section. Second, hsa-miR-629-3p strongly decreased steady-state TP53 levels while simultaneously inducing *CDKN1A*. These results indicate that the hits have diverse abilities to induce the nucleolar stress response for cell cycle interruption via upregulation of TP53 or *CDKN1A*.

### Two microRNA hits, hsa-miR-28-5p and hsa-miR-708-5p, are family members who each downregulate *RPS28*

During our mechanistic studies of the 15 microRNA hits, we hypothesized that the two included MIR-28 family members, hsa-miR-28-5p and hsa-miR-708-5p, would elicit similar results from each assay because of their identical seed sequences. These two microRNAs share the same 7 nt seed sequence (AGGAGCU) (**Figure 7A**) (52). Indeed, we observed similar cellular RB phenotypes following hsa-miR-28-5p or hsa-miR-708-5p treatment, including the same aberrant pre-rRNA processing signature and stark decreases in both mature 18S rRNA levels and global protein synthesis (**Figure 7A, Supplementary Table 6-7**). To uncover potential mechanisms for these phenotypic changes, we conducted RNAseq and differential expression analysis for cells treated with hsa-miR-28-5p or hsa-miR-708-5p mimics versus non-targeting siRNA (siNT) to control for nonspecific effects of small RNA transfection (**Figure 7B, Supplementary Table 11**). We hypothesized that differential gene expression should correlate strongly between the two microRNA siblings on a per-gene basis, given that the microRNAs should be largely sharing targets due to their identical seed sequences. Remarkably, when graphing per-gene log_2_ expression changes for hsa-miR-708-5p (y-axis) versus hsa-miR-28-5p (x-axis) relative to the negative control, the line of best fit was close to y = 0 + 1x with an *R*^2^ value of 0.61 (**Figure 7B**). Such a strong correlation indicates treatment with either hsa-miR-28-5p or hsa-miR-708-5p has a very similar effect on expression across the transcriptome, with the expression of individual genes increasing or decreasing, on average, to the same degree following treatment with either microRNA. These data strongly support the conclusion that treatment with hsa-miR-28-5p or hsa-miR-708-5p have similar phenotypic effects on MCF10A cells, and also that both microRNA hits cause highly-similar changes to the transcriptome as expected for two microRNAs with the same seed sequence.

**Figure 7.**
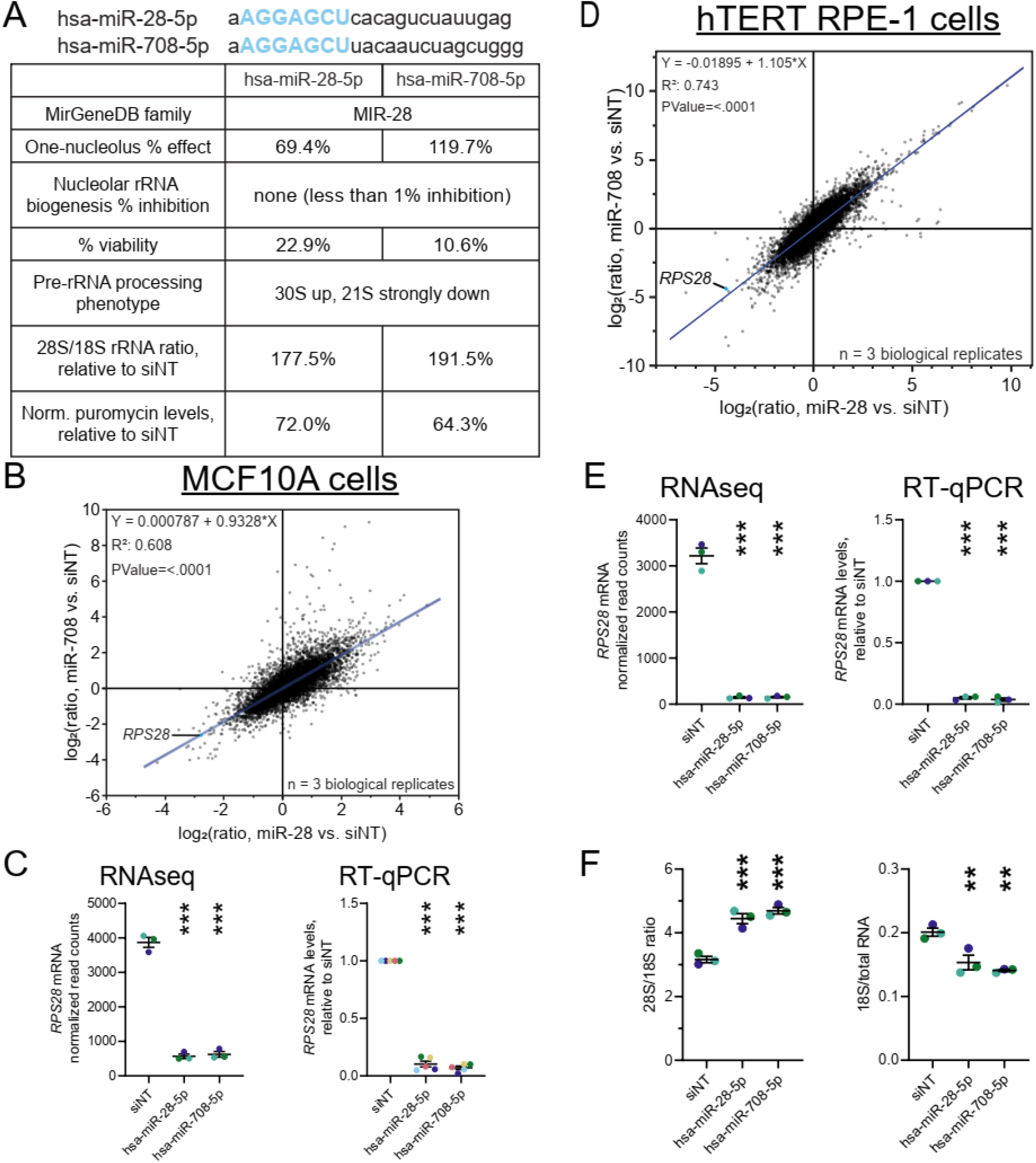
The MIR-28 siblings, hsa-miR-28-5p and hsa-miR-708-5p, elicit highly-similar transcriptomes in human MCF10A and hTERT RPE1 cells, including a potent reduction in *RPS28* levels. (A). Comparison of the MIR-28 family members, hsa-miR-28-5p and hsa-miR-708-5p. The full mature microRNA sequence is shown for each sibling, with the common AGGAGCU seed sequence indicated in blue. The table compares the RB phenotypes observed in MCF10A cells following treatment with either microRNA mimic. (B). Regression comparing log_2_-scale RNAseq differential expression profiles of MCF10A cells treated with either hsa-miR-28-5p or hsa-miR-708-5p, relative to siNT negative control. Each dot represents one mRNA, with *RPS28* labeled. The line-of-best-fit shown in blue, with equation, R^2^ value, and *p* value for non-zero slope indicated in top left. The data from 3 biological replicates were graphed and analyzed in JMP. (C). MIR-28-induced *RPS28* downregulation in MCF10A cells, measured by normalized RNAseq read counts or RT-qPCR analysis. For RNAseq reads, the mean ± SEM are shown alongside individual data points, colored by replicate (3 replicates). For RT-qPCR, data were normalized to 7SL RNA abundance as an internal control, then to siNT for comparison using the ΔΔC_T_ method. Mean ± SEM are shown alongside individual data points, colored by replicate (5 replicates). (D). Regression comparing log_2_-scale RNAseq differential expression profiles of hTERT RPE-1 cells treated with either hsa-miR-28-5p or hsa-miR-708-5p, relative to siNT negative control. Details are as written in (**B**). (E). MIR-28-induced *RPS28* downregulation in hTERT RPE-1 cells, measured by normalized RNAseq read counts or RT-qPCR analysis. Details are as written in (**C**). (F). Bioanalyzer analysis of total RNA isolated from control- or hit-treated hTERT RPE-1 cells. Left, the 28S/18S mature rRNA ratio; right, 18S mature rRNA/total RNA ratio. Mean ± SEM are shown alongside individual data points, colored by replicate. Data were graphed and analyzed by ordinary one-way ANOVA with multiple comparisons against siNT (non-targeting negative control) and Holm- Šídák correction in GraphPad Prism 8. **\***, *p* < 0.05; **\*\***, *p* < 0.01; **\*\*\***, *p* < 0.001.

Using our differential expression analysis, we followed up on *RPS28*, which was strongly downregulated by each MIR-28 sibling microRNA (**Figure 7B**). RPS28 is a ribosomal protein component of the 40S ribosomal subunit, and its depletion in human cells is reported to cause the same 30S up, 21S down aberrant pre-rRNA processing signature (101,102) that we observed following treatment with hsa-miR-28-5p or hsa-miR-708-5p. Indeed, in our own hands, siRNA-mediated *RPS28* knockdown reduces *RPS28* mRNA and RPS28 protein levels (**Supplementary Figure 6A-B**), decreasing nucleolar number and repressing nucleolar rRNA biogenesis in MCF10A cells (**Supplementary Figure 7A-C**). While RPS28 is not essential for maintaining normal 45S transcript levels, it may have a role in supporting RNAP1 promoter activity (**Supplementary Figure 7D-E**). We verified that RPS28 depletion causes the 30S pre-rRNA precursor to accumulate at the expense of 18S rRNA maturation (**Supplementary Figure 7F-H**), inhibiting global protein synthesis at a level commensurate with the depletion of the large ribosomal protein, RPL4 (**Supplementary Figure 7I**).

We validated the MIR-28-induced downregulation of *RPS28* by each microRNA sibling at the transcriptomic and protein levels in MCF10A cells (**Figure 7C, Supplementary Figure 8**). We also observed that, in hTERT RPE-1 cells, the MIR-28 siblings cause highly-similar transcriptomic changes and extremely potent downregulation of the *RPS28* mRNA in a dose-dependent manner (**Figure 7D-E, Supplementary Figure 9, Supplementary Table 12**). Furthermore, MIR-28 overexpression in hTERT RPE-1 also caused a stark decrease in mature 18S rRNA levels (**Figure 7F**), as observed in MCF10A cells (**Figure 5E**).

Given that both hsa-miR-28-5p and hsa-miR-708-5p treatment caused such a significant decrease in levels of *RPS28*, we hypothesized that this transcript may be directly targeted by the MIR-28 family. While the TargetScan (81), miRWalk (82), and miRDB (103) algorithms failed to predict such an interaction, DIANA microT-CDS (83) identified two tandem putative MIR-28 family binding sites in the *RPS28* mRNA 3’ UTR (**Supplementary Figure 10**). These tandem MIR-28 sites in *RPS28* are perfect 7mer seed matches conserved only within a subset of primates (**Supplementary Figure 10**); we hypothesize that this relatively low conservation throughout the animal kingdom may be the reason for false negatives using other prediction packages. By conducting *in silico* binding experiments using the DuplexFold algorithm on the RNAstructure server (84), we confirmed that the seed regions of both MIR-28 family members were predicted to favorably interact *in silico* with the candidate binding sites in *RPS28* (**Supplementary Figure 11A-D**). Scrambling the sequence of both binding sites in the candidate region strongly abrogated the predicted interaction with hsa-miR-28-5p or hsa-miR-708-5p (**Supplementary Figure 11E-H**).

We sought to test the extent to which the MIR-28 siblings could directly target these putative binding sites by using luciferase 3’ UTR reporter assays (**Figure 8A**) (86,104). Treatment with hsa-miR-28-5p or hsa-miR-708-5p inhibited reporter activity when the *Renilla* reporter contained the *RPS28* candidate region with WT putative MIR-28 binding sites (**Figure 8B**). Crucially, reporter assays containing the *RPS28* candidate region with scrambled putative binding sites completely rescued the microRNA-induced reporter downregulation (**Figure 8B**), supporting the conclusion that each MIR-28 family microRNA hit directly targets the *RPS28* 3’ UTR.

**Figure 8.**
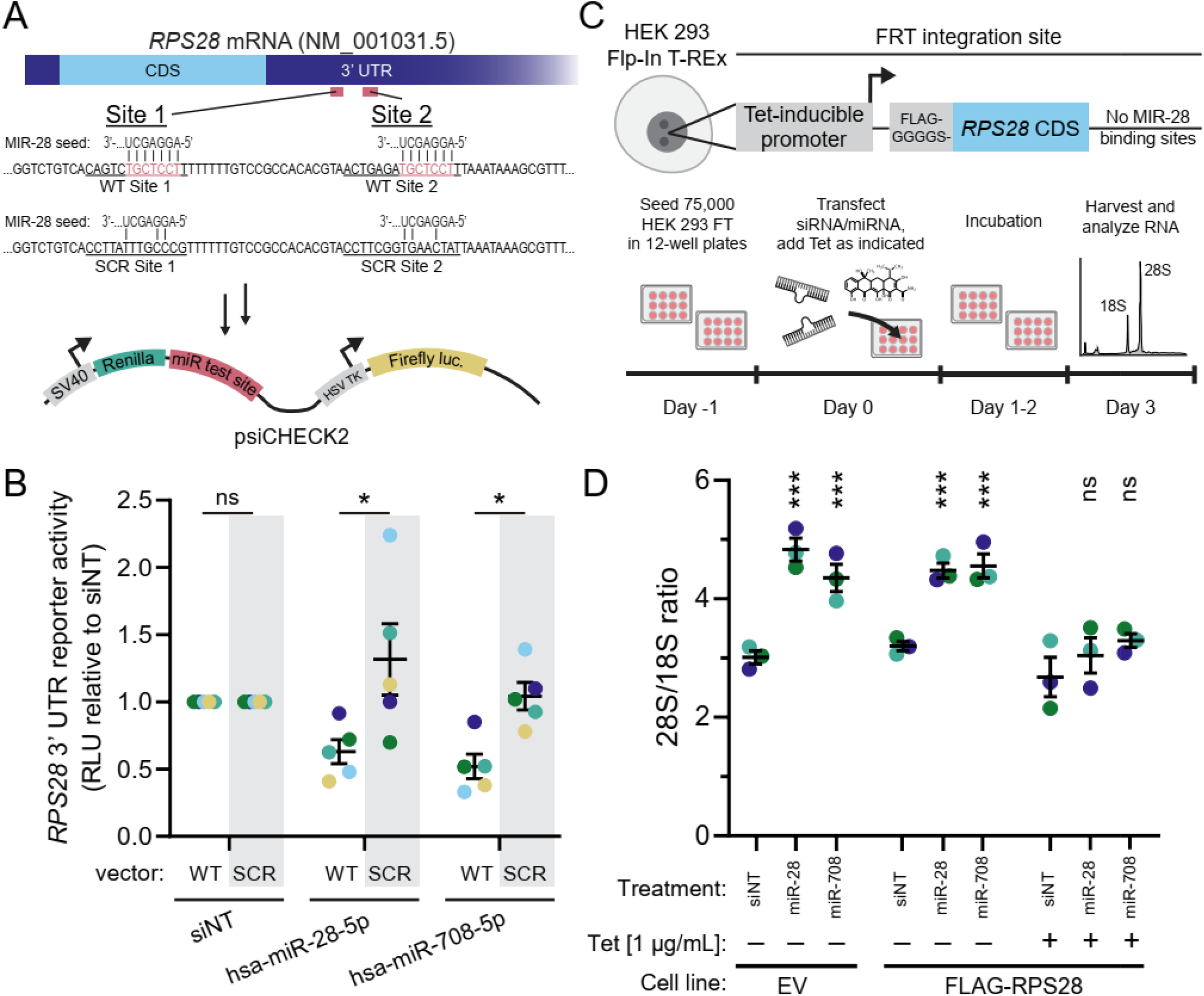
The MIR-28 siblings directly target tandem binding sites in the *RPS28* 3’ UTR to interrupt pre-18S processing. (A). Schematic for 3’ UTR luciferase reporter assay for testing direct binding of the MIR-28 sibl ngs to *RPS28*. Diagram for the *RPS28* mRNA transcript, indicating untranslated regions (UTR; dark blue) or coding sequence (CDS; light blue). Putative MIR-28 binding sites 1 and 2 are shown in red, and their wild-type (WT) sequence is given along with the design for scrambled MIR-28 sites (SCR) rescue construct. Antisense pairing of the 7mer MIR-28 seed is shown with the putative WT binding sites. WT or SCR regions of the *RPS28* 3’ UTR containing both putative MIR-28 sites were cloned into the miR test site in the 3’ UTR of the *Renilla* luciferase expression cassette in the psiCHECK2 plasmid. (B). Quantification of 3’ UTR luciferase reporter assays in (**A**), testing direct binding of MIR-28 siblings to *RPS28*. The vector construct is indicated as wild-type (WT) or scrambled (SCR) above the transfection treatment labels. Luciferase data were collected for each treatment and construct combination, then normalized to the siNT treatment on a per-construct basis. RLU stands for relative light units. Mean ± SEM are shown alongside individual data points, colored by replicate (5 replicates). The data were analyzed by unpaired two-sided Welch’s *t*-tests between WT and SCR constructs in GraphPad Prism 8. **\***, *p* < 0.05. (C). Schematic for rescue experiment conditionally expressing an *RPS28* mRNA lacking the MIR-28 family binding sites to recover normal rRNA maturation in the presence of MIR-28 sibling overexpression. (Top) Diagram of the engineered HEK 293 Flp-In T-REx cells containing a tetracycline-inducible FLAG-tagged *RPS28* cassette lacking the WT 3’ UTR harboring tandem MIR- 28 binding sites. (Bottom) Experimental outline indicating steps for cell seeding, simultaneous MIR-28 mimic transfection and engineered *RPS28* induction, incubation, and mature rRNA analysis by electropherogram. (D). Bioanalyzer analysis of the 28S/18S mature rRNA ratio from engineered HEK cells, treated as indicated. Treatment levels include siNT siRNA or hsa-miR-28-5p/hsa-miR-708-5p mimics. Induction with 1 μg/mL tetracycline (Tet) is indicated with a +. Cell line type was either the parental non- engineered line (empty vector, EV) or engineered line containing FLAG-tagged *RPS28* lacking WT 3’ UTR MIR-28 binding sites (FLAG-RPS28). Mean ± SEM are shown alongside individual data points, colored by replicate (*n* = 3 biological replicates). Data were graphed and analyzed by ordinary one- way ANOVA with multiple comparisons against siNT (non-targeting negative control) and Holm-Šídák correction in GraphPad Prism 8. ns, not significant; **\*\*\***, *p* < 0.001.

We also investigated if expression of a MIR-28-resistant *RPS28* could rescue normal pre-18S processing in the presence of the MIR-28 siblings. We genomically engineered HEK 293 Flp-In T-REx cells with a Tet-inducible FLAG-tagged *RPS28* cassette lacking the tandem MIR-28 binding sites normally present in the WT 3’ UTR (**Figure 8C**). We first verified that expression of the FLAG-tagged RPS28 protein was inducible only in the engineered cell line following treatment with 1 µg/mL tetracycline for 48 h (**Supplementary Figure 12**). Empty vector (EV) or non-induced *RPS28* HEK 293 Flp-In cells treated with either MIR-28 sibling showed an increase in the 28S/18S ratio versus siNT control, consistent with the defect in mature 18S rRNA production that we observed for these microRNA mimics in MCF10A cells (**Figure 8D and Figure 5E-F**). However, tetracycline-induced expression of the MIR-28-resistant *RPS28* construct restored normal 28S/18S ratios in the MIR-28 sibling overexpression background (**Figure 8D**). Coupled with the 3’ UTR reporter assays (**Figure 8 A-B)**, the rescue experiment provides extremely strong evidence that hsa-miR-28-5p and hsa-miR- 708-5p directly target *RPS28* to inhibit human pre-18S processing, a key step in RB (**Figure 9**).

**Figure 9.**
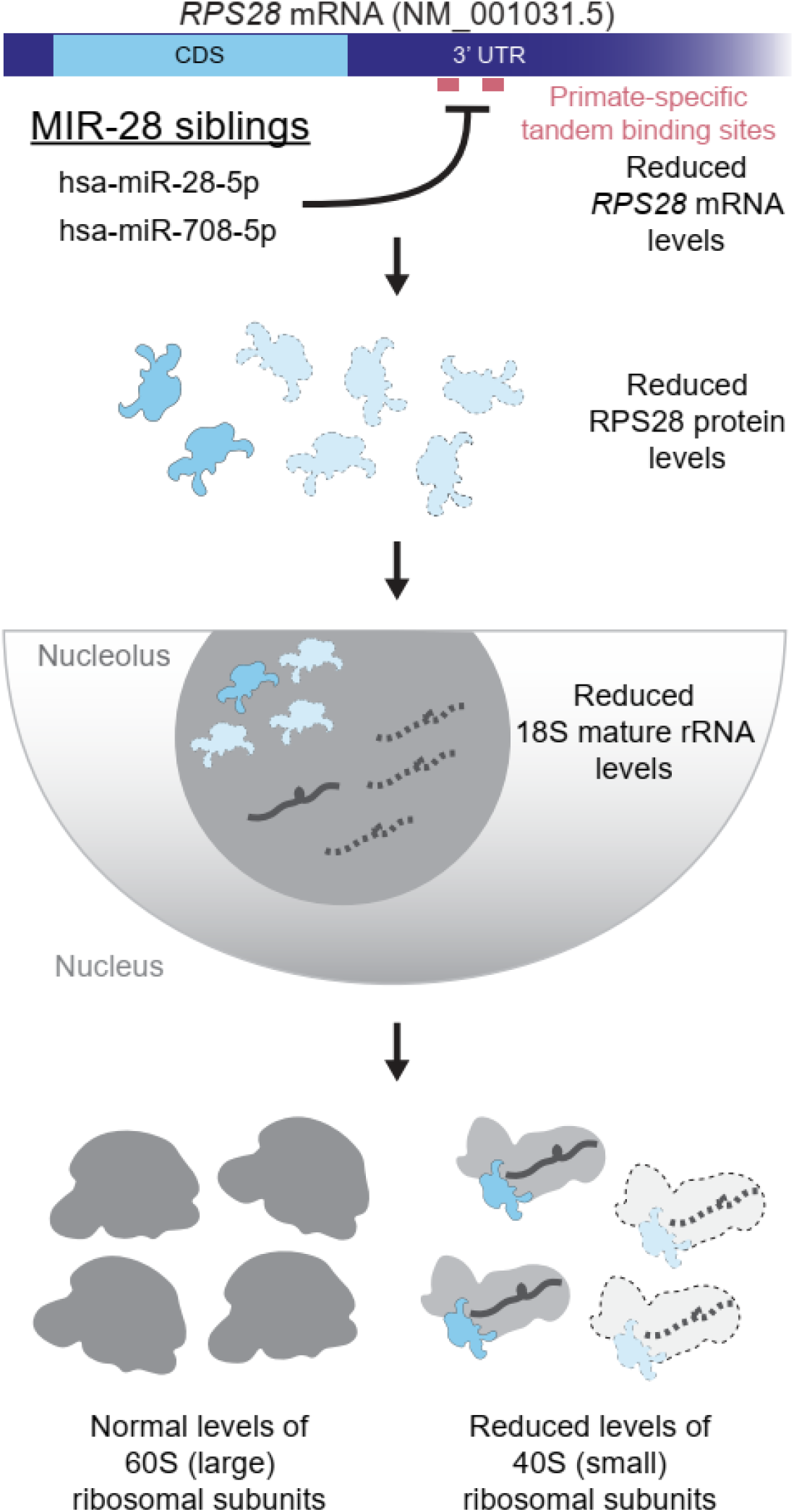
Model of the mechanism of MIR-28 sibling-mediated inhibition of ribosome biogenesis. The *RPS28* mRNA harbors two tandem, primate-specific binding sites for the MIR-28 microRNA family seed sequence. The MIR-28 siblings, hsa-miR-28-5p and hsa-miR-708-5p, can directly target *RPS28* at these sites, potently reducing *RPS28* mRNA and RPS28 protein levels. Lack of RPS28 protein in the nucleolus causes defects in pre-18S rRNA processing that lead to a reduction in levels of the mature 18S rRNA and subsequently 40S (small) ribosomal subunits.

To search for other possible direct targets of the MIR-28 family, we conducted miR-eCLIP analysis for MIR-28 microRNA:mRNA target chimeras (90). Briefly, treated cells were UV crosslinked, pooled, and an AGO2 immunoprecipitation was performed to isolate microRNAs actively complexed to their targets. Despite limit of detection challenges from modest AGO2 enrichment, sequencing revealed 9,243 high-confidence reads consisting of microRNA:mRNA chimeras representing functional target pairs (**Supplementary Table 13**). Each overexpressed MIR-28 sibling was loaded into AGO2 at levels commensurate with highly-expressed endogenous microRNAs (**Supplementary Figure 13A**), illustrating that the functional microRNAome was not expropriated by these transfected microRNA mimics. We uncovered 31 genes in our MIR-28 mimic RNAseq dataset bearing transcripts targeted by both MIR-28 siblings (**Supplementary Figure 13B**), including *MYC* and *CDKN1A* (*p21*) (**Supplementary Figure 14A-D**). An additional 113 genes in our MIR-28 mimic RNAseq dataset were targeted by either MIR-28 sibling (**Supplementary Figure 13**). Overall, direct MIR-28 targets identified by miR-eCLIP had reduced transcript levels following MIR-28 overexpression, as determined by RNAseq (**Supplementary Figure 13D-F**). Our data suggests that the MIR-28 siblings likely directly reduce *MYC* levels, providing another avenue of RB attenuation since MYC is an RNAP1 transcription factor (105). Targeting of *MYC* by the MIR-28 family has not previously been demonstrated in TarBase 8 (66). Furthermore, *CDKN1A* was directly targeted by the MIR-28 family, providing a logical explanation for the lack of change of *CDKN1A* mRNA levels in the face of TP53 upregulation that we observed (**Figure 6D-E**, hsa-miR-28-5p or hsa-miR-708-5p). While *RPS28* was not observed to be a direct target of hsa-miR-28-5p or hsa-miR-708-5p by miR-eCLIP, this negative result may be the product of subpar detection limits. Together, our findings strongly support the ability of the MIR-28 family to perturb the transcriptome with a high degree of similarity among human cell lines, and to engage in direct post-transcriptional downregulation of the central RB regulator *MYC* and of *RPS28*, an RP critical for normal 18S rRNA maturation.

## Discussion

Our work represents the first systematic venture into uncovering the complex roles of microRNAs as governors of ribosome biogenesis. Using our unbiased high-content screening platform for changes in nucleolar number, we have uncovered 72 novel microRNA negative regulators of RB. Strikingly, 51/72 hits strongly inhibited nucleolar rRNA biogenesis as measured by nucleolar 5-EU incorporation, supporting a role for the hits in antagonizing RB. We highlight 27 validated, conserved microRNA hits present in MirGeneDB, including 17 hits that induce strong repression of nucleolar rRNA biogenesis in multiple human cell lines. Stringent selection and mechanistic validation of a subset of 15 novel microRNA hits unexpectedly revealed a major effect of hit overexpression to be dysregulation of 30S pre-rRNA processing. Prior to our work, no specific microRNAs had yet been observed to directly affect pre-rRNA processing (35). While hits in the subset did not appear to reliably alter RNAP1 transcription, almost all subset hits inhibited global protein synthesis and caused upregulation of *CDKN1A* (*p21*), with nearly half increasing TP53 steady-state levels. We hypothesized that the microRNA hits were acting by binding to mRNAs required for nucleolar function and reducing their levels. Bioinformatics of all the microRNA hits revealed that they were enriched for mRNA targets encoding proteins localized within the nucleolus or bearing functions in cell cycle progression, proliferation, or TP53 signaling, supporting our mechanistic hypothesis.

To further probe the hypothesis that microRNA mimic hits preferentially target nucleolar proteins, we focused on two MIR-28 family members, hsa-miR-28-5p and hsa-miR-708-5p. We chose them because they share the same 7 nt AGGAGCU seed sequence and their overexpression causes the same RB defects, including a severe pre-18S rRNA processing defect. Comparison of RNAseq results following overexpression of either mimic resulted in highly-similar transcriptomic profiles in two distinct non-cancerous human cell lines, MCF10A (breast epithelial) and hTERT RPE-1 (retinal pigment epithelium), leading us to focus on *RPS28* as a putative direct target. Using established experimental methods to assess direct targeting of microRNAs, we found that these two MIR-28 family members directly target the *RPS28* 3’ UTR, providing a convincing explanation for the observed pre-18S rRNA processing defect (101,102) and supporting our model (**Figure 9**). Critically, we showed that conditional co-expression of the MIR-28 siblings with an *RPS28* mRNA lacking MIR- 28 binding sites in engineered HEK 293 cells was able to restore normal proportions of the 28S and 18S rRNAs, verifying that the targeting of *RPS28* by the MIR-28 family members is the mechanism that underlies the RB defect. Taken together, our microRNA mimic screen has revealed that the MIR- 28 family directly targets the mRNA encoding the ribosomal protein RPS28, leading to a drastic reduction in its levels with concomitant defects in RB and a decrease in cell viability in human cells.

Our unexpected discovery that the *RPS28* 3’ UTR harbors primate-specific MIR-28 binding sites underscores the importance of conducting unbiased functional experiments in human systems to advance basic and translational biology. MIR-28 targeting of *RPS28* had not previously been predicted or observed [reviewed in (41)], which is surprising given the extensive published literature on this microRNA family and the ubiquitous presence of the essential RPS28 protein in all growing cells. The 2023 version of DIANA microT-CDS predicts the MIR-28 family binding sites in the 3’ UTR of human *RPS28*, though other common microRNA target prediction algorithms including TargetScan, miRDB, and miRWalk fail to predict these interactions. Furthermore, the tandem MIR-28 binding sites in the *RPS28* 3’ UTR can be found only in the most related of primates, including humans, old world monkeys, and apes. We hypothesize that this lack of conservation may be the reason that previous experiments in model systems failed to identify these interactions despite strong conservation of RB and the MIR-28 family in animals. Our findings make a strong argument for undertaking unbiased screens in order to discover novel microRNA targets in human cells, instead of relying solely on *in silico* predictions or non-human model organisms.

While our results point to the novel microRNA hits canonically acting to inhibit RB by post-transcriptionally downregulating target genes with nucleolar localization or functions in the cell cycle, we cannot exclude the possibility that these microRNAs may also have a more immediate role inside the nucleolus itself. A number of studies have defined nucleolar subsets of microRNAs in mammalian cells (38–40,106,107), though the function of nucleolar microRNAs remains poorly understood. It has been suggested that the nucleolus may serve as a staging platform for microRNAs to complex with target mRNAs outside the competitive, mRNA-rich cytoplasm (107), or perhaps that efflux of microRNAs from the nucleolus may be part of a stress response to the invasion of foreign genetic material (40). Additionally, AGO2 has been observed to bind to regions of rDNA possibly via rRNA-mediated tethering (37), though more research is needed to fully understand the potential of microRNAs to directly downregulate the 45S transcript. Previous studies (38,40) have observed the nucleolus to contain a number of our hits’ families, including miR-19b, miR-25, miR-34a, miR-182, miR-183, miR-192, miR-330, and miR-629. However, in most cases, the microRNA strand was not indicated in these studies. Future investigation of nucleolar microRNAs may shed additional light on the potential direct impacts of the hits inside the nucleolus beyond the present scope.

Based on the promiscuous nature of microRNA activity, screens with microRNA mimics have an increased scale and complexity of direct regulatory perturbation compared to previous RB screening campaigns (53,54,108–110). While prior screens used siRNA technology to surgically deplete expression of a single gene, the transfection of a microRNA mimic may directly deplete tens to thousands of mRNAs. Simultaneous manipulation of multiple gene regulatory networks with microRNAs may lead to complex, potentially-discordant cellular phenotypes. This is because assay results report an integration of many more heterogeneous expression changes than in simpler single-gene siRNA experiments. One key example from this work is that the MIR-28 siblings elevate TP53 protein levels without a concomitant increase in TP53’s downstream target, *CDKN1A* (*p21*), as expected (111). Our miR-eCLIP data indicated that both MIR-28 microRNAs in fact directly target *CDKN1A in vivo*, as previously shown for hsa-miR-28-5p (112), resolving this discrepancy. While we found a straightforward explanation for this case, other discrepancies in our data likely remain. These results invite additional probing to improve our understanding of the complex functional perturbations associated with each microRNA hit.

Looking forward, we highlight our discovery of novel microRNA negative regulators of RB in the context of cancer. Differential regulation of many of the 72 hits has been observed in various tumors (94,113–146), with hits including hsa-miR-28-5p and hsa-miR-708-5p (147–156) often acting as tumor suppressors. For example, hsa-miR-28-5p overexpression impedes growth of colorectal cancer cell lines and mice tumor xenografts (152) as well as prostate cancer cell lines (156). We propose that targeting of *RPS28* by members of the MIR-28 family may be one novel aspect of their effectiveness as tumor suppressors. We emphasize the enrichment of the hits’ targets for involvement in the cell cycle, which is tightly intertwined with RB, nucleolar formation, and tumorigenesis (25,54,157). Additionally, microRNA mimic therapeutics are a promising avenue in oncology (158), though roadblocks including delivery and dosage have impeded achieving success in the clinic (159). Combining mimics and small molecules may enable inhibition of multiple targets or decrease required microRNA dose (159), as seen in clinical trials for hepatitis C (160,161). Given that cancer cells often hyperactivate ribosome production (24), our study underscores the potential of harnessing conserved microRNAs for chemotherapy as standalone therapeutics or in concert with other potent small molecule inhibitors of RB like BMH-21 to simultaneously target pre-rRNA transcription and processing.

## Acknowledgements and Funding Sources

We thank the members of the laboratory of S.J.B. and S. Carter for insightful information, questions, and comments throughout the manuscript writing process. We thank P. Pawlica and J. Steitz for advice on the microRNA 3’ UTR luciferase assays and for sharing the psiCHECK2 plasmid (86). We thank the Yale Center for Genome Analysis (YCGA) for performing Agilent Bioanalyzer analysis, RNAseq library preparation and sequencing. We acknowledge the use of CellProfiler for image analysis [(56), http://www.cellprofiler.org/]. This work was supported by the following grants from the National Institutes of Health (NIH): 1R35GM131687 (to S.J.B.), 1F31DE030332 (to M.A.M.), and T32GM007223 (to C.J.B., M.A.M., and S.J.B.).

---

**Supplementary Figure 1.**
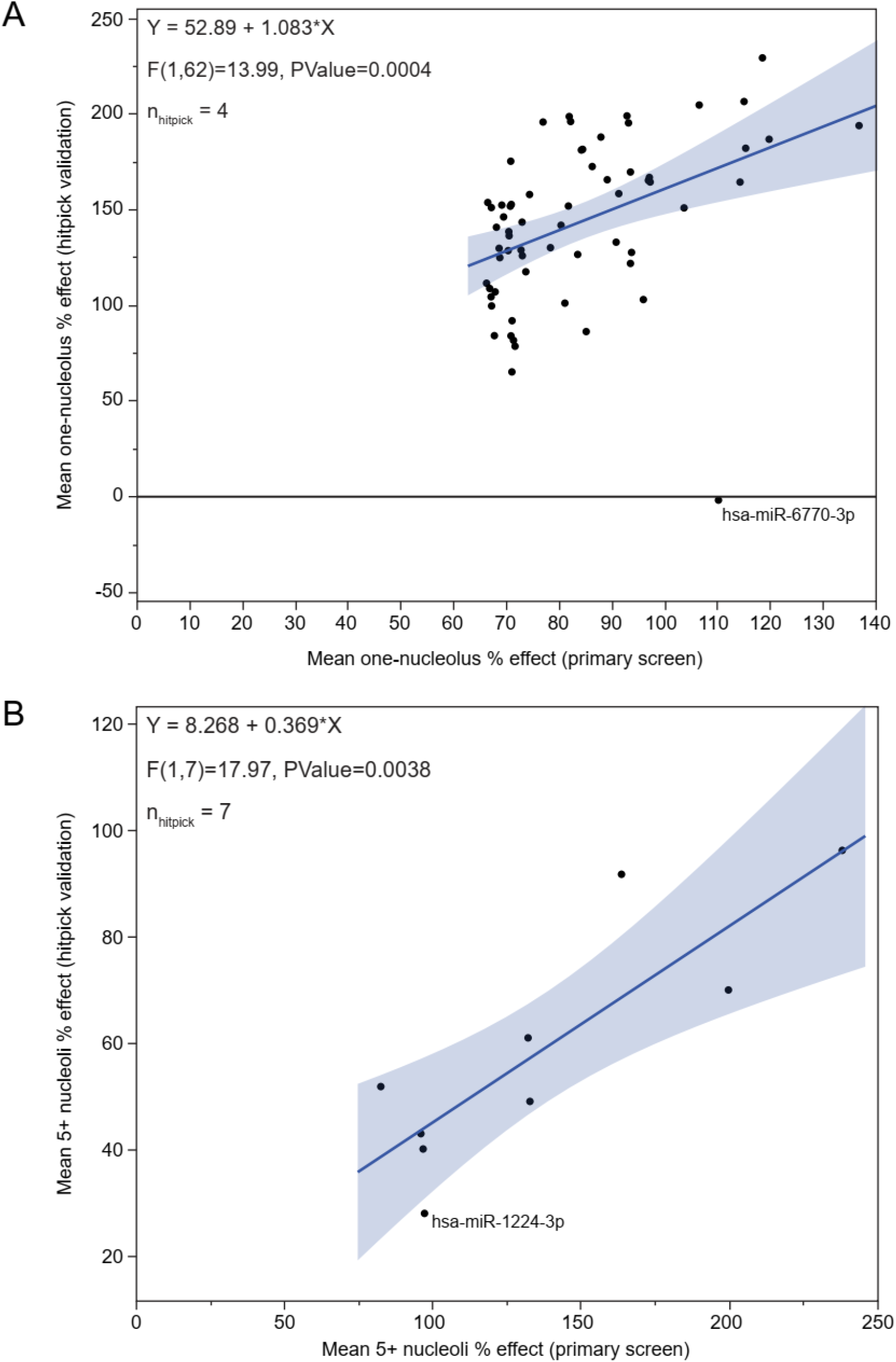
Primary screen hitpick validation excludes hsa-miR-6770-3p and hsa-miR-1224-3p. (A) Comparison of mean one-nucleolus percent effects from primary screen (x-axis) or hitpick validation (y-axis) for 64 one-nucleolus hits. The one-nucleolus hitpick was conducted with *n* = 4 replicates. Graphing and linear regression was performed in JMP. hsa-miR-6770-3p was excluded from downstream analysis. (B) Comparison of mean 5+ nucleoli percent effects from primary screen (x-axis) or hitpick validation (y-axis) for nine 5+ nucleoli hits. The 5+ nucleoli hitpick was conducted with *n* = 7 replicates. Graphing and linear regression was performed in JMP. hsa-miR-1224-3p was excluded from downstream analysis.

**Supplementary Figure 2.**
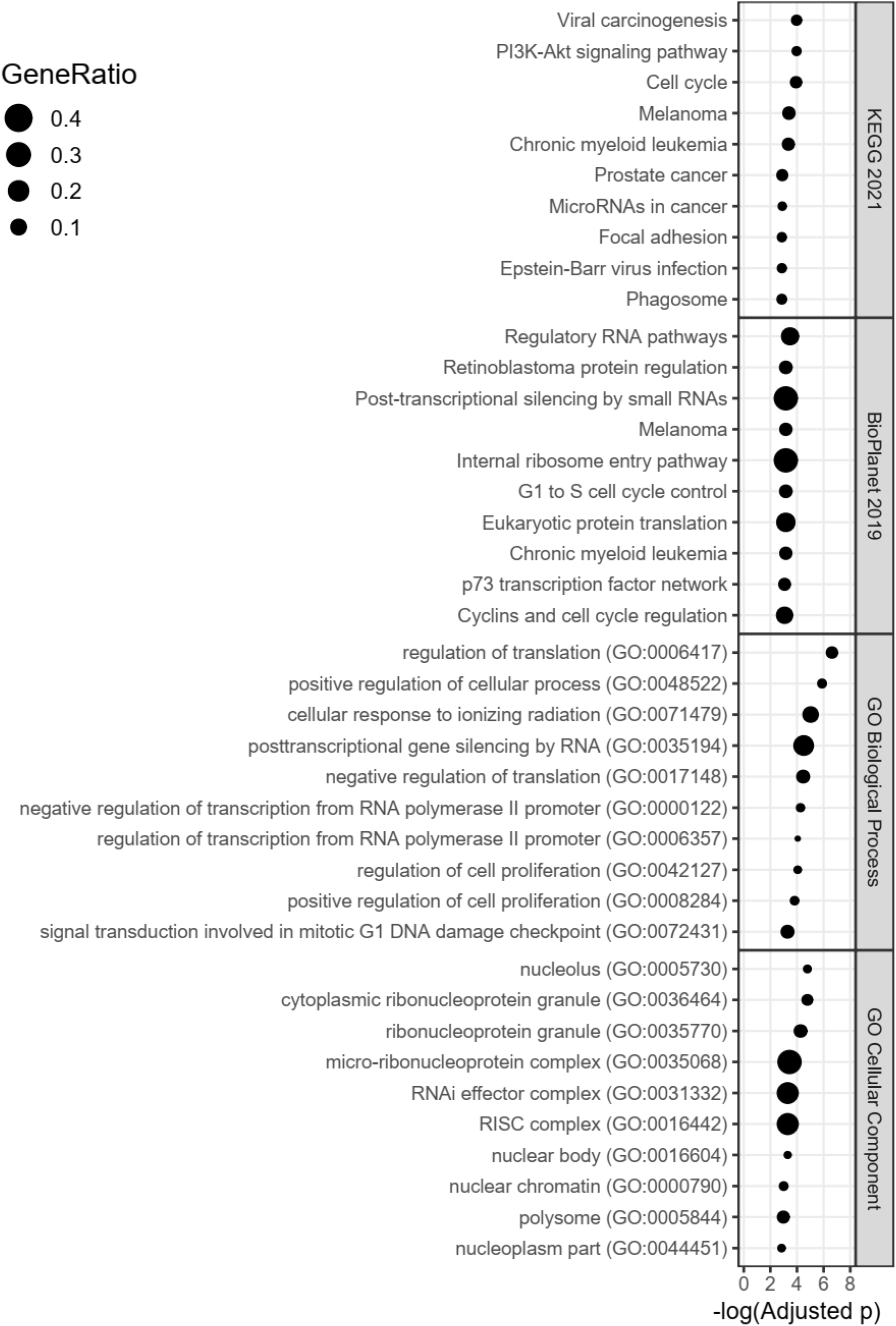
Targets of 24 MirGeneDB hits represented in TarBase 8 are enriched for regulators of RNA pathways, translation, the cell cycle, and for localization in the nucleolus. Enrichment plots for 135 genes targeted by 5 or more of the microRNA hits present in MirGeneDB. Plots indicate −log_10_(adjusted *p*) on the x-axis and the gene ratio as the marker size. Enrichment analysis was conducted with Enrichr, and plots were made in R. Enrichment databases: Kyoto Encyclopedia of Genes and Genomes (KEGG) 2021; NCATS BioPlanet of Pathways 2019; Gene Ontology (GO) Biological Process 2018; GO Cellular Component 2018.

**Supplementary Figure 3.**
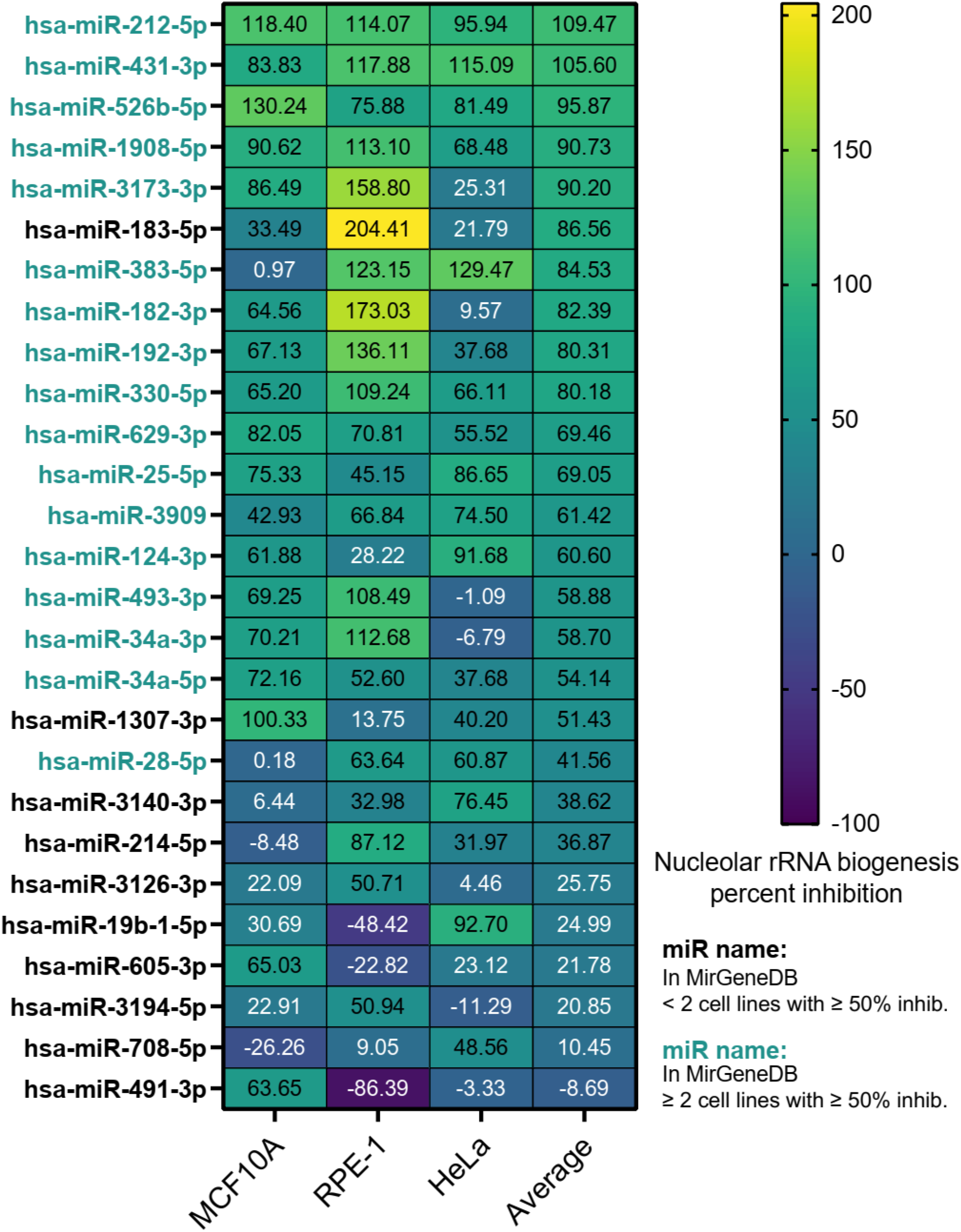
Conservation of nucleolar rRNA biogenesis inhibition by MirGeneDB microRNA hits across 3 human cell lines. Heatmap showing the nucleolar rRNA biogenesis percent inhibition following overexpression of 27 microRNA mimic hits present in MirGeneDB in 3 diverse human cell lines. Cell types are indicated on the x-axis, with an unweighted average column also calculated. All microRNA hit names are bolded because each microRNA is present in MirGeneDB. MicroRNA hit names are colored depending on conservation across cell lines. Hits whose overexpression causes at least a 50% inhibition of nucleolar rRNA biogenesis in two or more cell lines (teal name); hits failing this criterion (black name).

**Supplementary Figure 4.**
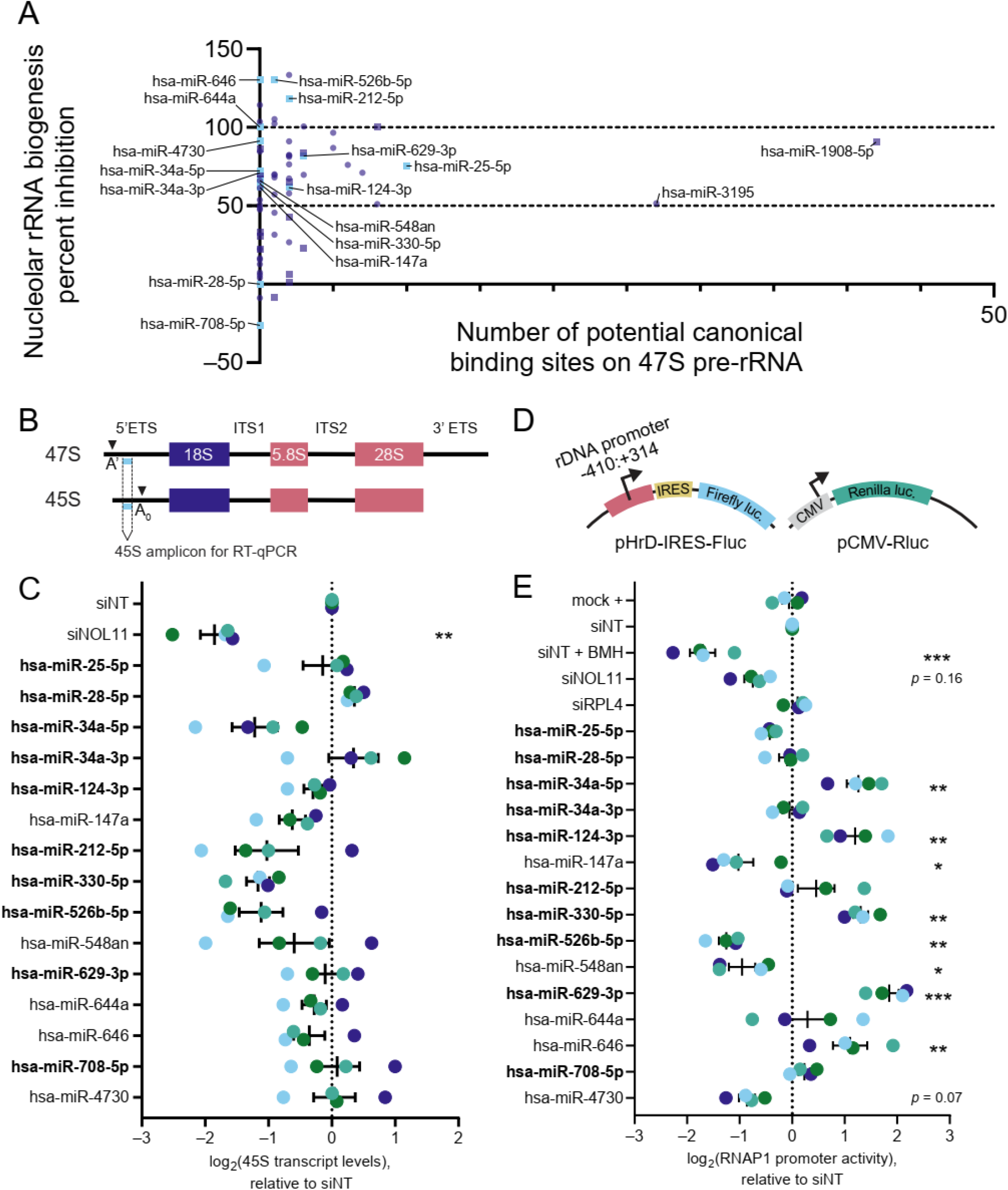
MicroRNA hits do not reliably alter RNAP1 transcription. (A). Scatter plot comparing the number of potential canonical (seed) binding sites on the 47S pre-rRNA transcript to the nucleolar rRNA biogenesis percent inhibition for the 72 microRNA hits, as predicted by BLAST. A number of hits have close to 0 predicted canonical 47S binding sites but strong nucleolar rRNA biogenesis percent inhibition. Select hits are labeled, and labels for percent inhibition are shown at 50% inhibition (empirical assay cutoff, consistent with pre-rRNA modification defect) and 100% inhibition (siPOLR1A positive control, consistent with pre-rRNA transcription defect) (55). The data were graphed in GraphPad Prism 8. (B). Schematic indicating the amplicon (light blue) used for 45S pre-rRNA RT-qPCR, located between the A’ and A_0_ cleavage sites in the 5’ ETS of the primary rRNA transcript. The location of the mature 18S, 5S and 28S rRNAs are indicated. (C). RT-qPCR analysis of levels of the 45S pre-rRNA precursor transcript as a proxy for RNAP1 transcription. Hit names are bolded for MirGeneDB members. The mean ± SEM are shown alongside individual data points, colored by replicate (4 replicates). The data were normalized to 7SL RNA abundance as an internal control, then to siNT for comparison using the ΔΔC_T_ method. The data were analyzed by ordinary one-way ANOVA with multiple comparisons against siNT and Holm-Šídák correction in GraphPad Prism 8. **\***, *p* < 0.05; **\*\***, *p* < 0.01; **\*\*\***, *p* < 0.001. (D). Schematic for the dual-luciferase reporter assay for RNAP1 promoter activity (54,99). MCF10A cells were co-transfected with pHrD-IRES-Fluc, in which a 724 bp fragment of the rDNA promoter and early 5’ ETS drive firefly luciferase production, and pCMV-Rluc, in which the constitutive CMV promoter drives *Renilla* luciferase production. (E). Dual-luciferase reporter assay for RNAP1 promoter activity. Hit names are bolded for MirGeneDB members. The mean ± SEM are shown alongside individual data points, colored by replicate (4 replicates). The data were analyzed by ordinary one-way ANOVA with multiple comparisons against siNT and Holm-Šídák correction in GraphPad Prism 8. **\***, *p* < 0.05; **\*\***, *p* < 0.01; **\*\*\***, *p* < 0.001.

**Supplementary Figure 5.**
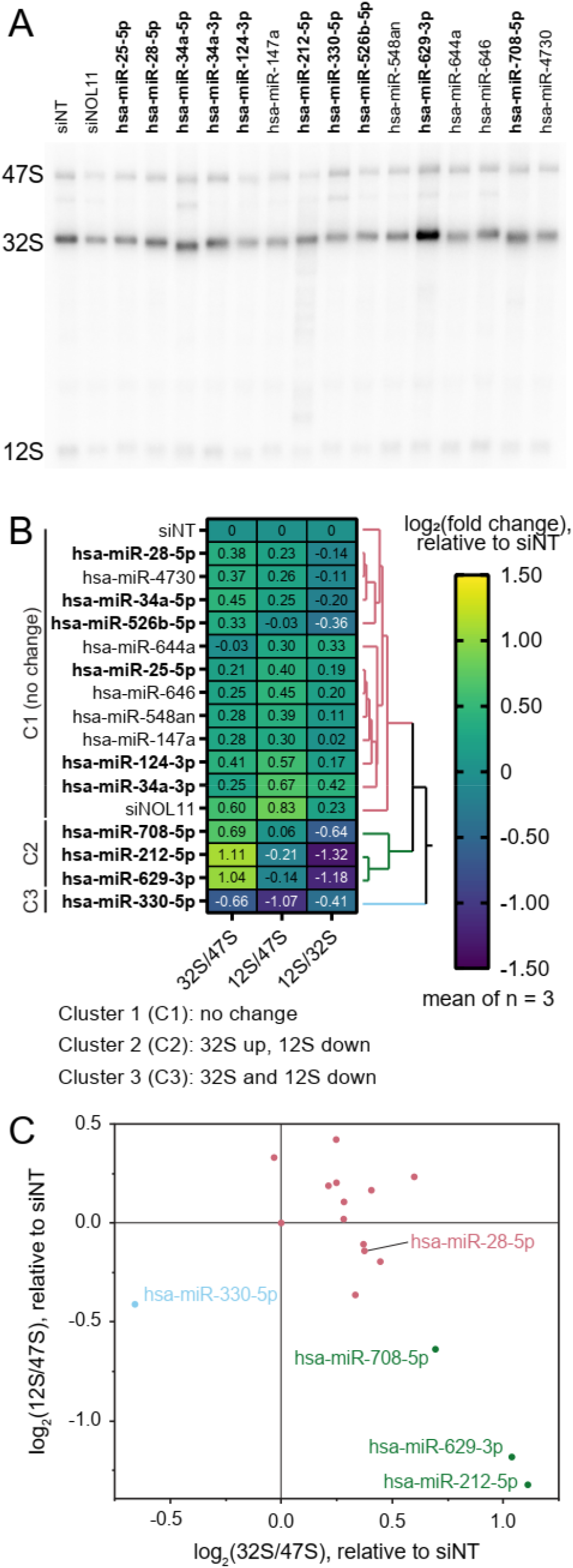
Four microRNA hits slightly interfere with ITS2 processing. (A). Representative ITS2 (probe P4) northern blot of 3 μg of total RNA isolated from control- or hit-treated MCF10A cells. The pre-rRNA processing intermediates are labeled on the left according to the schematic in Figure 5A. The images were quantified using Bio-Rad Image Lab. Hit names are bolded for MirGeneDB microRNAs. (B). Clustered heatmap showing log_2_-transformed Ratio Analysis of Multiple Precursor [RAMP, (77)] calculations for microRNA mimic hits, normalized to si non-targeting (siNT) negative control. Values represent mean log_2_-scale RAMP ratio for *n* = 3 replicates. Clusters: no change (C1, red); 32S up, 12S down (C2, green); both 32S and 12S down (C3, blue). The RAMP ratios were calculated in Microsoft Excel. Three clusters were assigned using hierarchical Ward clustering in JMP, and data were graphed in GraphPad Prism 8. Hit names are bolded for MirGeneDB microRNAs. (C). Log_2_-scale 32S/47S and 12S/47S pre-rRNA precursor RAMP ratios after subset microRNA mimic treatment, relative to a non-targeting (siNT). Cluster colors the same as in (**B**). The data were graphed in GraphPad Prism 8, with select hits labeled.

**Supplementary Figure 6.**
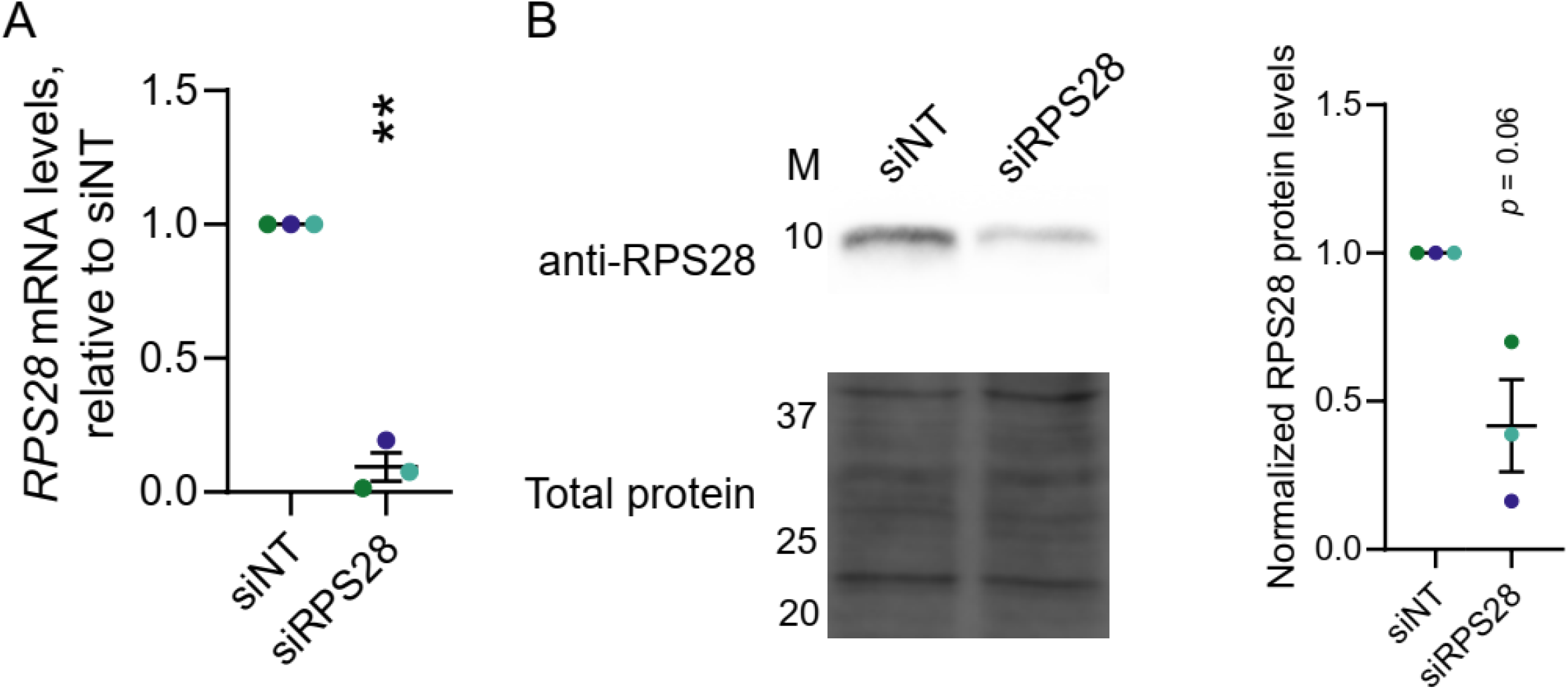
Validation of siRNA-mediated knockdown of RPS28. (A) RT-qPCR quantification of mRNA transcript levels following siRNA knockdown of *RPS28* in MCF10A cells. The data were normalized to 7SL RNA abundance as an internal control, then to a non-targeting siRNA (siNT) for comparison using the ΔΔC_T_ method. The mean ± SEM are shown alongside individual data points, colored by replicate (3 replicates). Data were analyzed by unpaired two-sided Welch’s *t*-tests in GraphPad Prism 8. **\***, *p* < 0.05; **\*\***, *p* < 0.01; **\*\*\***, *p* < 0.001. (B) Immunoblot quantification of protein levels following siRNA knockdown for *RPS28* in MCF10A cells. Anti-RPS28 blotting is shown in the example immunoblot. Total protein is the trichloroethanol total protein stain loading control. M, molecular weight marker lane in kDa. The images were quantified using Bio-Rad Image Lab. The mean ± SEM are shown alongside individual data points, colored by replicate (3 replicates). The data were graphed and analyzed by unpaired two-sided Welch’s *t*-tests in GraphPad Prism 8. **\***, *p* < 0.05; **\*\***, *p* < 0.01; **\*\*\***, *p* < 0.001.

**Supplementary Figure 7.**
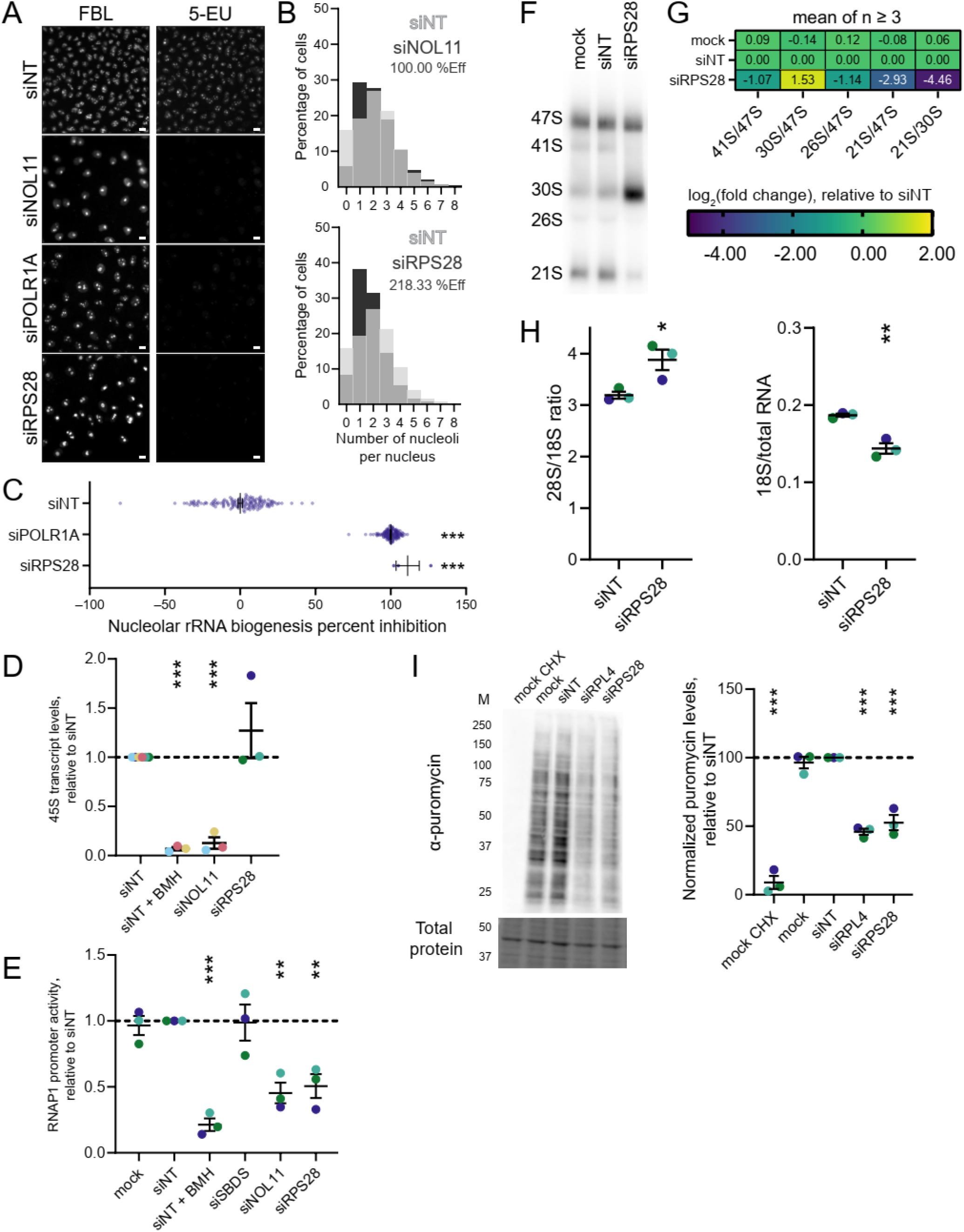
RPS28 knockdown decreases nucleolar number, inhibits nucleolar rRNA biogenesis, interdicts pre-18S processing, and reduces global protein synthesis. (**A**). Representative images of control-treated or RPS28-depleted MCF10A cells following fibrillarin (FBL) antibody staining and 5-EU incorporation. FBL immunostaining and 5-EU click labeling are shown as separate channels. Scale bars, 10 μm. siNT is a non-targeting negative control siRNA. siNOL11, NOL11 knockdown is a positive control for the one-nucleolus phenotype. siPOLR1A, POLR1A (RPA194) knockdown is a positive control for nucleolar rRNA biogenesis inhibition. siRPS28, RPS28 knockdown. (**B**) Quantification of nucleolar number following siRNA depletion of RPS28. The histograms indicate the percentage of cells with a given number of nucleoli. The siNT negative control histogram is shown in light gray on all graphs for reference. Histograms from cells depleted of NOL11 or RPS28 are shown in black. siNOL11 is the positive control for decrease in nucleolar number. The one-nucleolus percent effect for black-labeled treatment is shown. (C). Quantification of the percent inhibition of nucleolar rRNA biogenesis. The overall mean percent inhibition ± SEM is shown for each treatment, with each dot representing one well. The siNT negative control is set to 0% inhibition, and the siPOLR1A positive control is set to 100% inhibition. Data were analyzed by ordinary one-way ANOVA with multiple comparisons against siNT and Holm-Šídák correction in GraphPad Prism 8. **\*\*\*** *p* < 0.001. (D). RT-qPCR analysis of levels of the 45S pre-rRNA as a proxy for RNAP1 transcription. siNT, non-targeting siRNA negative control. siNT + BMH, siNT transfection plus acute 1 µM BMH-21 pre-treatment to selectively inhibit RNAP1 activity. siNOL11, NOL11 depletion as a positive control. siRPS28, RPS28 depletion. The mean ± SEM are shown alongside individual data points, colored by replicate (3 replicates). The data were normalized to 7SL RNA abundance as an internal control, then to siNT for comparison using the ΔΔC_T_ method. Data were analyzed by ordinary one-way ANOVA with multiple comparisons against siNT and Holm-Šídák correction in GraphPad Prism 8. **\*\*\***, *p* < 0.001. (E). Dual-luciferase reporter assay for RNAP1 promoter activity (54,99). Mock, no siRNA mock transfection control. siNT, non-targeting siRNA negative control. siNT + BMH, siNT transfection plus acute 1 µM BMH-21 pre-treatment to selectively inhibit RNAP1 activity. siSBDS, depletion of the cytoplasmic pre-60S assembly factor SBDS as a negative control. siNOL11, NOL11 depletion as a positive control. siRPS28, RPS28 depletion. The mean ± SEM are shown alongside individual data points, colored by replicate (3 replicates). The data were analyzed by ordinary one-way ANOVA with multiple comparisons against siNT and Holm-Šídák correction in GraphPad Prism 8. **\*\***, *p* < 0.01; **\*\*\***, *p* < 0.001. (F). Representative ITS1 (probe P3) northern blot of 3 μg of total RNA isolated from RPS28-depleted MCF10A cells. Mock, no siRNA mock transfection control. siNT, non-targeting siRNA negative control. siRPS28, RPS28 depletion. The pre-rRNA processing intermediates are labeled on the left. The images were quantified using Bio-Rad Image Lab. (G). Heatmap showing log_2_-transformed Ratio Analysis of Multiple Precursor [RAMP (77)] calculations, normalized to the si non-targeting (siNT) negative control. The values represent mean RAMP ratio for *n* = 4 replicates, except *n* = 3 for mock. RAMP ratios were calculated in Microsoft Excel and data were graphed in GraphPad Prism 8. (H). Bioanalyzer analysis for 1 μg of total RNA isolated from RPS28-depleted MCF10A cells. Left, the 28S/18S mature rRNA ratio; right, the 18S mature rRNA/total RNA ratio. Mean ± SEM are shown alongside individual data points, colored by replicate (3 replicates). The data were graphed and analyzed by ordinary one-way ANOVA with multiple comparisons against a non-targeting siRNA (siNT) and Holm-Šídák correction in GraphPad Prism 8. **\***, *p* < 0.05; **\*\***, *p* < 0.01. (I). Representative examples of SUnSET puromycin incorporation assay (53,80) immunoblots of total protein isolated from RPS28-depleted MCF10A cells. α-puromycin is an example blot for puromycin incorporation as a proxy for global protein synthesis. Total protein is the trichloroethanol total protein stain loading control. M, molecular weight marker in kDa. Mock, no siRNA mock transfection control. Mock CHX is mock cells co-treated for 1 h with puromycin and 100 µg/mL cycloheximide to halt protein translation. siNT, non-targeting siRNA negative control. siRPL4, depletion of the 60S component RPL4 as a positive control. siRPS28, RPS28 depletion. Images were quantified with Bio-Rad Image Lab. The mean ± SEM are shown alongside individual data points, colored by replicate (3 replicates). Data were normalized to a non-targeting siRNA (siNT), then graphed and analyzed by ordinary one-way ANOVA with multiple comparisons against siNT and Holm-Šídák correction in GraphPad Prism 8. **\*\*\***, *p* < 0.001.

**Supplementary Figure 8.**
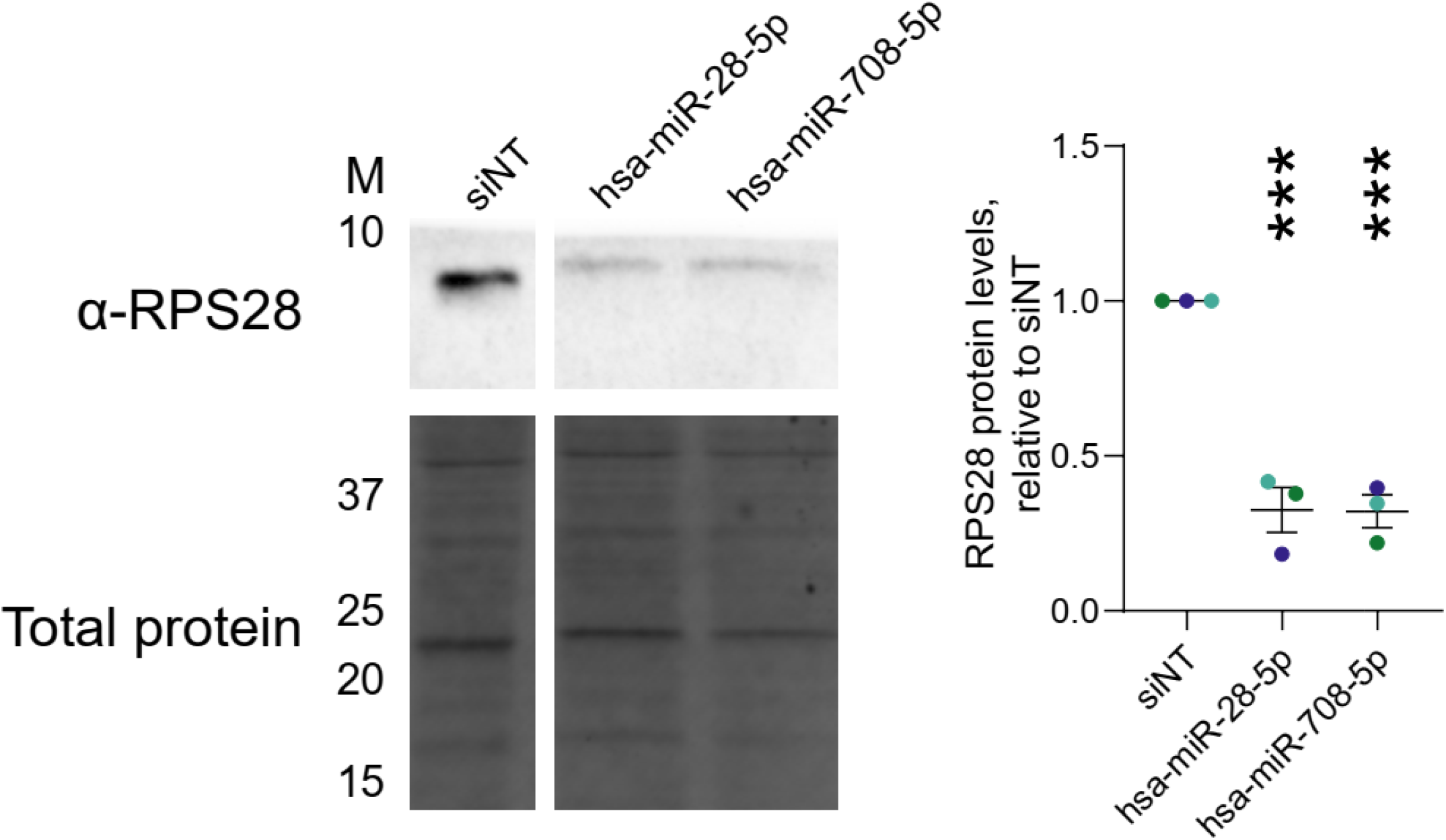
RPS28 protein levels are potently reduced by the MIR-28 siblings in MCF10A cells. Immunoblot analysis and quantification of RPS28 protein levels from control- or MIR-28 sibling- treated MCF10A cells. α-RPS28 indicates the example immunoblot for the labeled protein. Total protein, trichloroethanol total protein stain loading control. M, molecular weight marker lane in kDa. siNT, non-targeting siRNA negative control. hsa-miR-28-5p or hsa-miR-708-5p, transfected MIR-28 microRNA mimics. The images were quantified with Bio-Rad Image Lab. Mean ± SEM are shown alongside individual data points, colored by replicate (3 replicates). The data were normalized to siNT, then graphed and analyzed by ordinary one-way ANOVA with multiple comparisons against siNT and Holm-Šídák correction in GraphPad Prism 8**\*\*\***, *p* < 0.001.

**Supplementary Figure 9.**
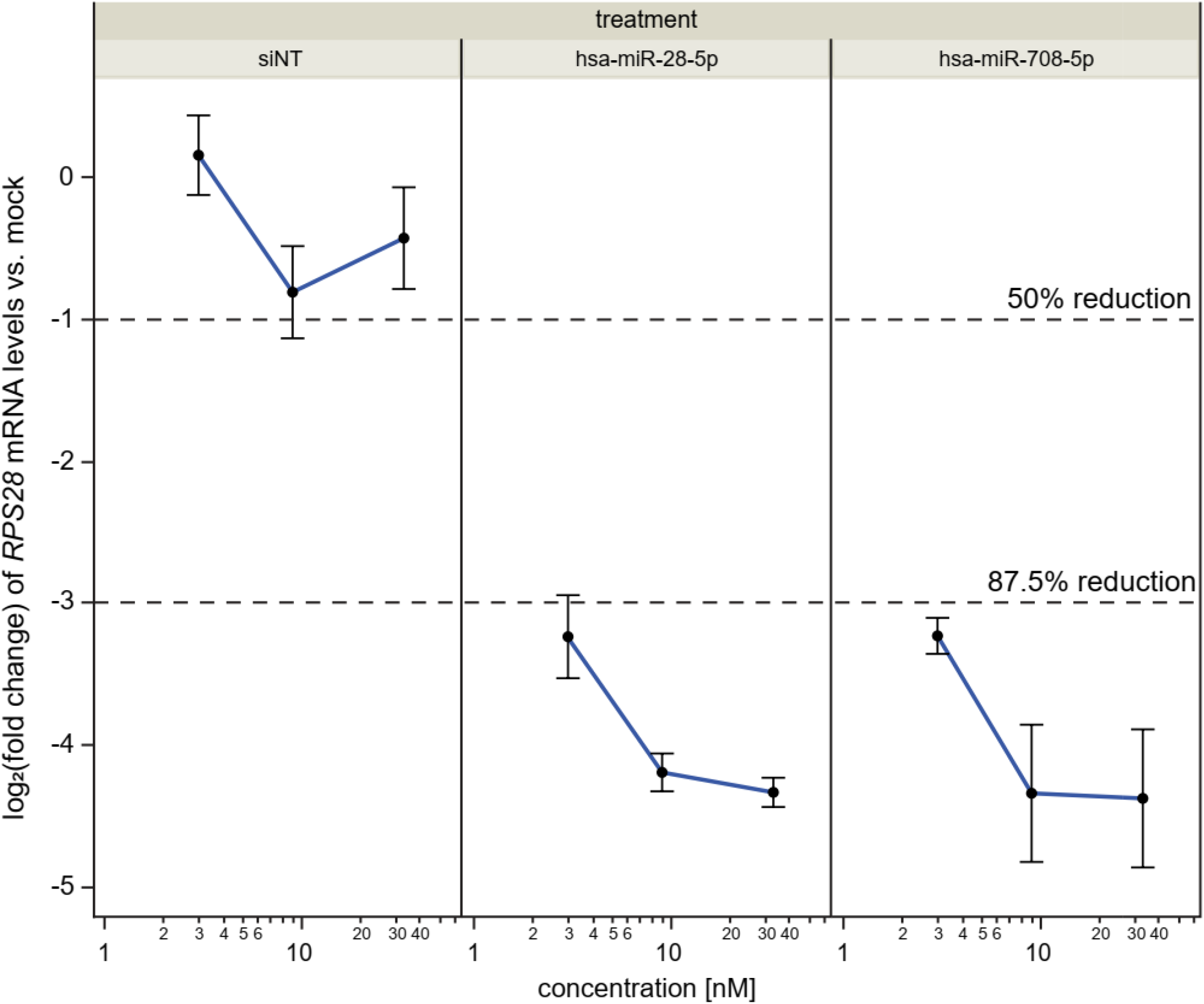
*RPS28* mRNA levels are potently inhibited by the MIR-28 siblings in hTERT RPE-1 cells. Three-point concentration response curves quantifying the MIR-28 siblings’ effect on *RPS28* mRNA levels in hTERT RPE-1 cells by RT-qPCR. siNT, non-targeting siRNA negative control. Hsa-miR-28- 5p or hsa-miR-708-5p, transfected MIR-28 microRNA mimics. Concentrations of siRNA/microRNA mimics were 33 nM, 9 nM, and 3 nM. Log_2_ fold change of *RPS28* mRNA levels compared to mock transfection is shown on the y-axis. Dashed lines indicate the relative linear fold change associated with -1 or -3 log_2_ units. The data were normalized to 7SL RNA abundance as an internal control, then to mock for comparison using the ΔΔC_T_ method. Data are graphed as mean ± SEM for *n* = 3 biological replicates and *n* = 2 technical replicates.

**Supplementary Figure 10.**
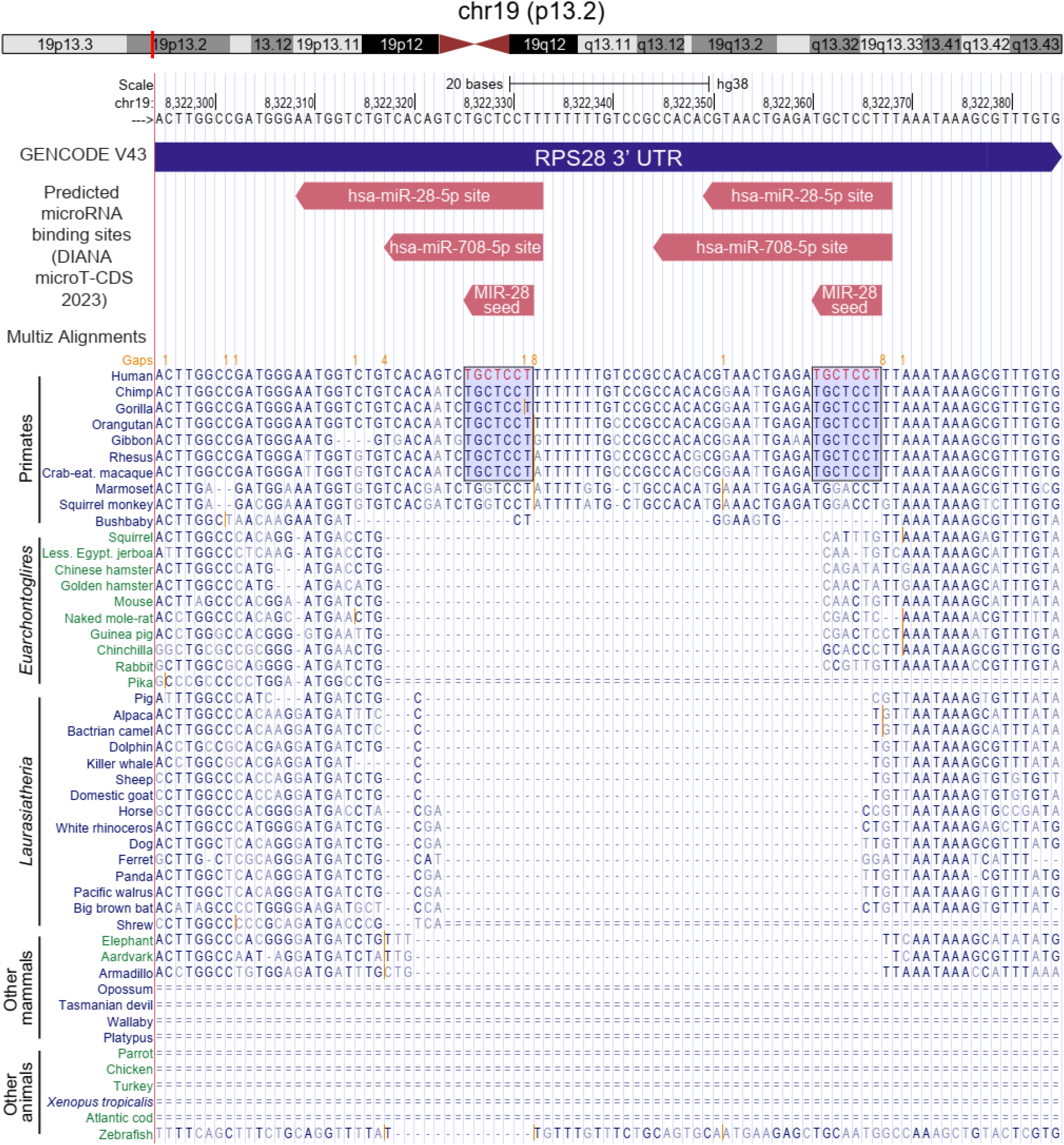
The predicted tandem MIR-28 family binding sites in the RPS28 3’ UTR are conserved only within a subset of primates. UCSC Genome Browser visualization of the *RPS28* 3’ UTR region containing predicted tandem MIR- 28 family binding sites. A schematic of the locus on human chromosome 19 is shown above the Browser view. A DIANA microT-CDS 2023 track with predicted binding sites and the MIR-28 family seed sequence is shown in red. Multi-species genomic alignments for selected species in the Multiz 100 Vertebrate Species Alignment and Conservation track are shown, with group labels annotated on the left. Conservation of the predicted 7mer MIR-28 sites within a subset of primates is highlighted with a light blue box.

**Supplementary Figure 11.**
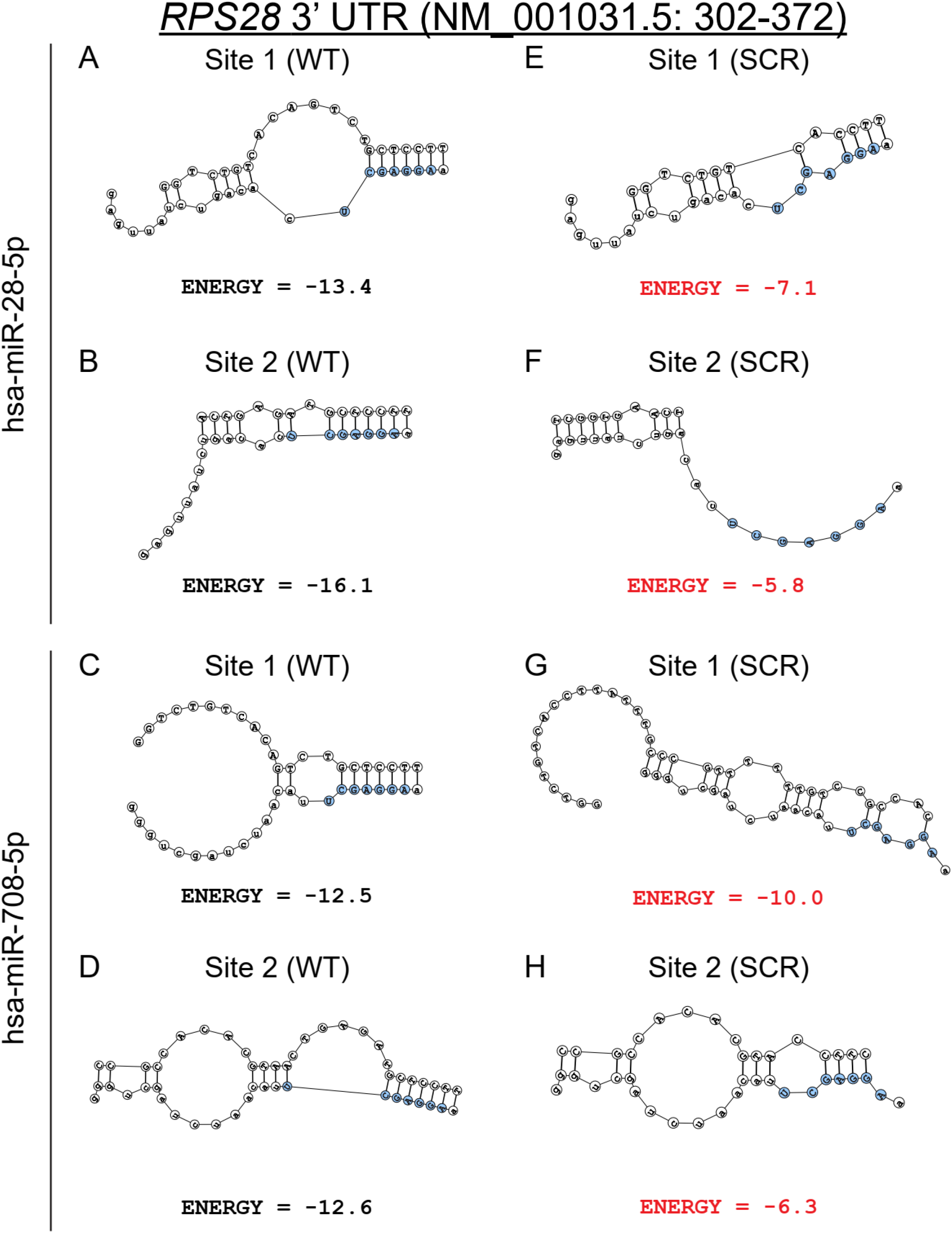
Predicted binding structures between MIR-28 siblings and putative binding sites in the *RPS28* 3’ UTR. (**A-D**). DuplexFold predicted structures between hsa-miR-28-5p (**A-B**) or hsa-miR-708-5p (**C-D**) to wild-type (WT) putative MIR-28 sites in the *RPS28* 3’ untranslated region (UTR). The seed sequence for each microRNA is shown in blue. The DuplexFold predicted binding energies are indicated for each structure. Transcriptomic coordinates for the binding region input are shown in the header. Extraneous target base pairs were removed for visualization. (**E-H**). Corresponding DuplexFold predicted structures between hsa-miR-28-5p (**E-F**) or hsa-miR-708- 5p (**G-H**) to seed site scrambled (SCR) putative MIR-28 sites in *RPS28* 3’ untranslated region (UTR). Each binding energy (red) is reduced following seed scrambling. Extraneous target base pairs were removed for visualization.

**Supplementary Figure 12.**
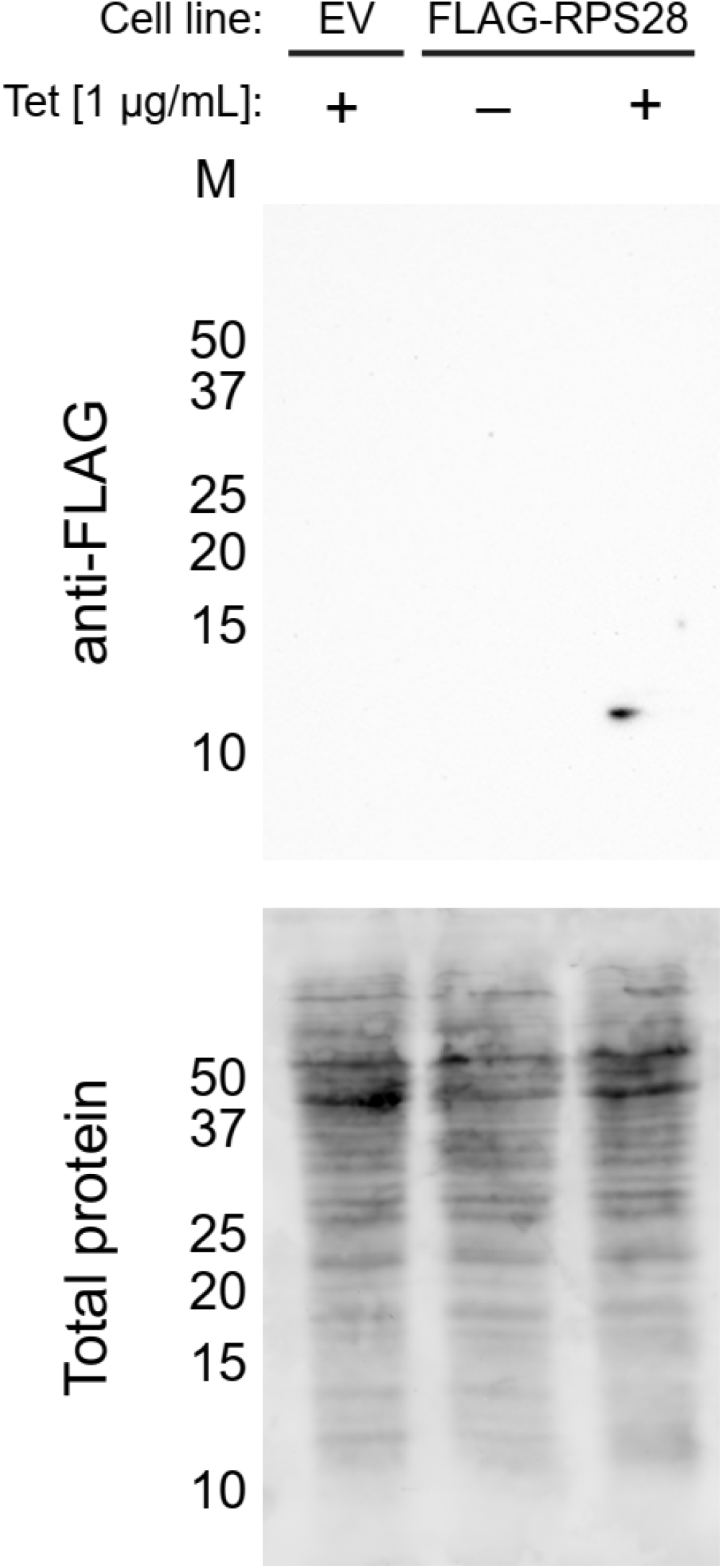
Tetracycline-inducible expression of FLAG-tagged RPS28 in an engineered HEK 293 Flp-In T-REx cell line. Immunoblot analysis of FLAG-tagged RPS28 protein levels from HEK 293 Flp-In T-REx cells. Anti-FLAG indicates an immunoblot for the FLAG tag. Total protein, trichloroethanol total protein stain loading control. M, molecular weight marker lane in kDa. Cell line, parental non-engineered/empty vector (EV) or FLAG-RPS28 engineered HEK 293 Flp-In cells. Tet, tetracycline induction of protein expression at 1 µg/mL for 48 h. +, positive tetracycline induction; –, no tetracycline induction.

**Supplementary Figure 13.**
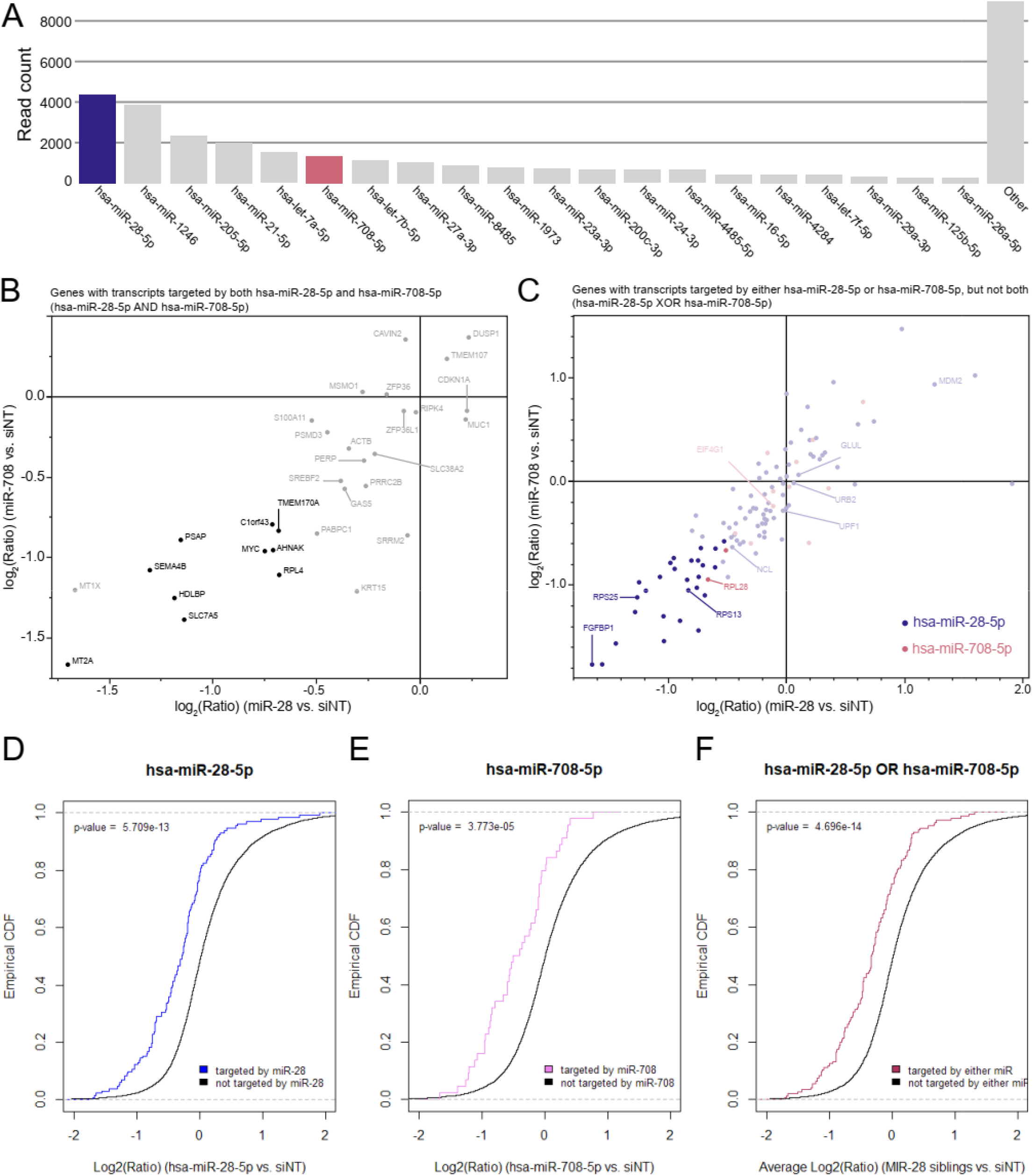
Direct targets of the MIR-28 siblings revealed by miR-eCLIP. (A). Read counts of microRNAs sequences enriched following MIR-28 mimic overexpression and miR-eCLIP AGO2 biochemical enrichment. Transfected mimics of the MIR-28 siblings hsa-miR-28-5p (blue) and hsa-miR-708-5p (red) have similar read counts compared to other endogenous microRNAs (gray). (B). Plot of genes with mRNA transcripts targeted by both hsa-miR-28-5p and hsa-miR-708-5p. The miR-eCLIP data were filtered for genes expressed in MCF10A cells, and RNAseq differential expression data following MIR-28 treatment is graphed for each gene on the x- and y-axes. Bolded points represent genes downregulated by both MIR-28 mimics by at least -0.5 log_2_ units, with a differential expression false discovery rate < 0.05 for both treatments. All genes are labeled with their HGNC symbol. (C). Plot of genes with mRNA transcripts targeted exclusively (XOR) by either hsa-miR-28-5p or hsa-miR-708-5p. The miR-eCLIP data were filtered and graphed as above. The genes encoding mRNA transcripts were bound by hsa-miR-28-5p (blue) or bound by hsa-miR-708-5p (red), according to miR-eCLIP data. Select genes are labeled with HGNC symbol. (**D-F**). Cumulative empirical distribution plots indicating RNAseq differential expression data for labeled groups of targets. Genes not targeted by MIR-28 siblings (black) are compared to genes targeted by hsa-miR-28-5p (**D**, blue line), by hsa-miR-708-5p (**E**, pink line), or by either MIR-28 sibling (**F**, maroon line) based on miR-eCLIP results. Data were analyzed by Kolmogorov-Smirnov testing in R.

**Supplementary Figure 14.**
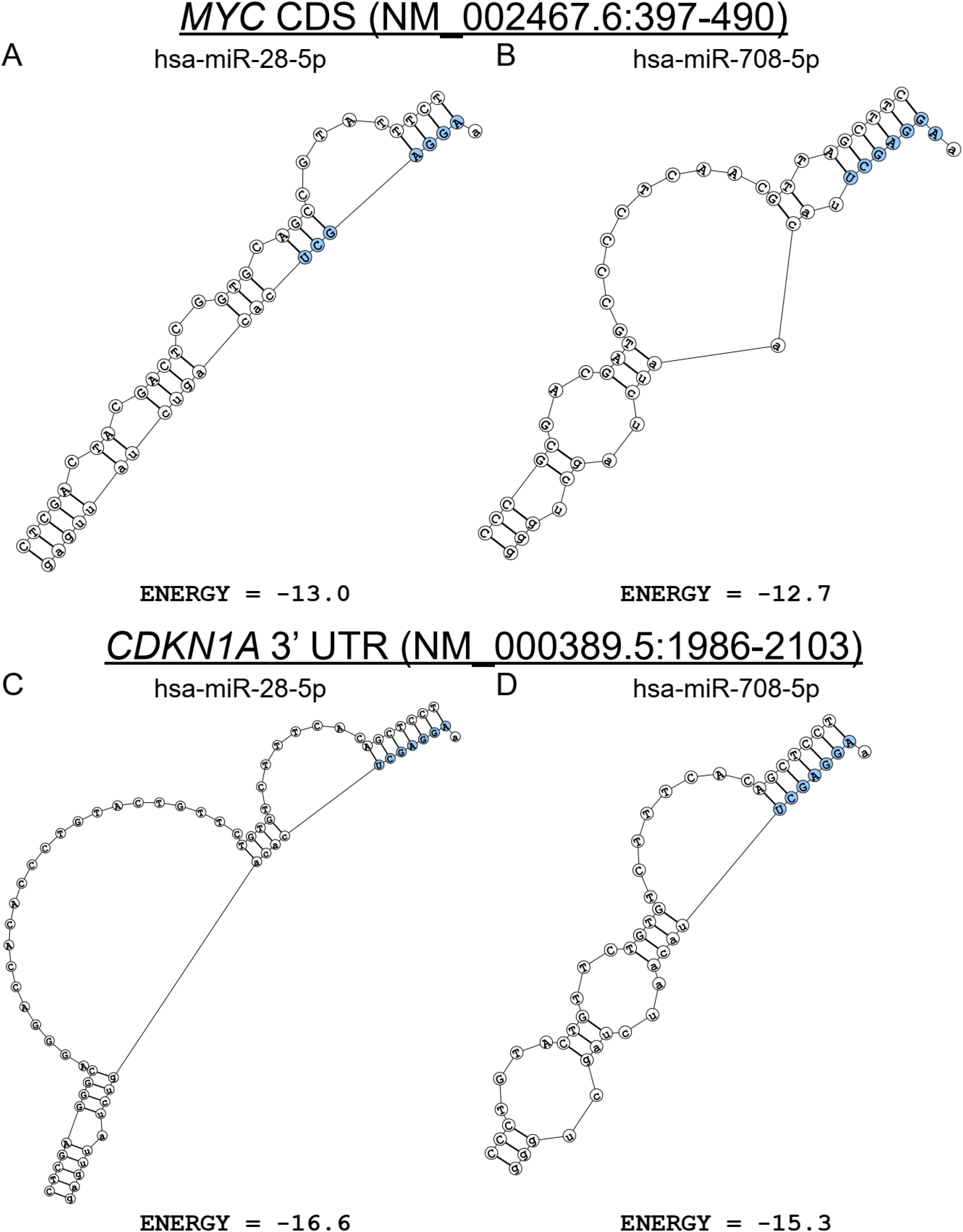
Predicted MIR-28 binding structures in the *MYC* CDS or *CDKN1A* (*p21*) 3’ UTR. (**A-B**). DuplexFold predicted structures between hsa-miR-28-5p (**A**) or hsa-miR-708-5p (**B**) and wild-type (WT) putative MIR-28 sites in the *MYC* coding sequence (CDS). The seed sequence for each microRNA is shown in blue. The DuplexFold predicted binding energies are indicated for each structure. Transcriptomic coordinates for the binding region input are shown in the header. Extraneous target base pairs were removed for visualization. (**C-D**). DuplexFold predicted structures between hsa-miR-28-5p (**C**) or hsa-miR-708-5p (**D**) and wild-type (WT) putative MIR-28 sites in *CDKN1A* (*p21*) 3’ UTR. Other features as in (**A-B**).

